# Genetic innovation in coronaviruses driven by a viral nuclease

**DOI:** 10.64898/2026.08.15.744938

**Authors:** Christopher Bianco, Alex C. Stabell, Manivel Lodha, Miranda Aldis, Michael A Tartell, Theodora Hatziioanou, Paul D. Bieniasz

## Abstract

Genetic variation in viruses is well known to arise from polymerase-driven nucleotide misincorporation. However, insertion and deletion (indel) mutations that occur at lower, largely unknown, frequencies can underly more dramatic phenotypic changes that emerge when advantageous. Using a human coronavirus (HCoV-OC43) construct that reports rare indel mutations, we show that non-structural protein-15 (NSP15), a nuclease encoded by coronaviruses, can drive the acquisition of a class of insertion mutations. Ultra-deep sequencing of both HCoV-OC43 and SARS-CoV-2 populations reveals a similar requirement for NSP15 during insertion mutant generation. Overall, the insertional mutation frequency exceeded 10^-3^/genome in these two coronaviruses. Analysis of thousands of HCoV-OC43 and SARS-CoV-2 insertion mutants reveals a mutational process in which NSP15 cuts viral RNA, yielding oligonucleotides that correspond to inserts that are acquired at distal genomic locations. The presence of an insertion mutation at the S1/S2 junction in the SARS-CoV-2 spike protein that generates a furin cleavage site and enhances viral transmissibility, may have been necessary for enabling the COVID19 pandemic. We found numerous examples of potential furin cleavage site acquisition and replacement through insertion mutation during the normal course of coronavirus replication. Such events are, therefore, likely commonplace in coronavirus populations of a size that occurs in nature.

## Main

Genetic variation in viruses enables phenotypic changes, including evasion of immune defenses and colonization of new hosts. While polymerase mediated nucleotide misincorporation is a key driver of evolutionary change in RNA viruses^1,2^, profound phenotypic changes, such as acquisition of new functions, are generally accompanied by more substantial genetic changes, such as insertion mutations. The frequency with which insertion mutations are generated in viruses is normally below that measurable in most assays^1,3^, and insertions are typically revealed only when an associated selective advantage elevates their frequency in a viral population. Moreover, the comparative rareness with which insertion mutations are encountered renders the molecular mechanisms underlying their occurrence mysterious^1,3,4^.

Insertion mutations in RNA viruses can be highly consequential. The most notorious example is an apparent 12 nucleotide insertion in the spike (S) gene of SARS-CoV-2 relative to its likely sarbecovirus ancestors^5,6^. This insertion generates a site recognized by furin-like proteases at the S1/S2 boundary, enabling spike protein cleavage during virion genesis. This furin cleavage site (FCS) can enhance spike fusogenicity ^7,8^, but may also reduce trimer stability, and thus has context dependent effects on viral fitness^7,9,10^. In some animal models, the FCS can increase transmissibility or pathogenicity^10,11^ and it is possible that FCS acquisition by a SARS-CoV-2 progenitor was necessary for the COVID19 pandemic.

The NSP15 protein, also termed endonuclease U (Endo-U), is a ribonuclease encoded by all known coronaviruses, with homologs also found in the broader nidovirus order^12^. Through its nuclease activity, which acts immediately 3’ to U and sometimes C nucleotides^13,14^, NSP15 mitigates the innate immune response to coronaviruses by cleaving the viral genome and thus reducing the level of viral double stranded (ds)RNA^13,15–18^. NSP15 was also reported to increase levels of subgenomic RNA and reduce levels of RNA species with deletions^19^. Herein, we deploy two approaches, based on bioassays of viral populations and viral sequencing at extreme depth, to document a previously undescribed mutagenic function for coronavirus NSP15 proteins. We show that NSP15 drives the generation of a class of insertion mutations in the genomes of two coronaviruses (HCoV-OC43 and SARS-CoV-2) during the normal course of their replication. As part of these studies, we show that insertion mutations are encountered at a frequency in excess of 10^-3^/genome in viral populations expanded over a few generations from a molecularly cloned founder. These findings lead to the conclusion that naturally occurring HCoV-OC43 and SARS-CoV-2 populations of a size often found within a single host likely contain millions of insertion mutants, many generated through the action of NSP15.

## Results

### An HCoV-OC43 indicator virus that reports indel mutations

Using a human coronavirus molecular clone, we built an indicator construct that would report the occurrence of insertion or deletion (indel) mutations during virus replication (HCoV-OC43/fsGFP(WT)) (**Fig. 1a**). This viral genome encoded a frameshifted (fs) GFP sequence, in place of the non-essential gene *ns2*. Specifically, a 34 nucleotide (11 codons +1 nucleotide) leader sequence, inserted between the methionine initiator codon and the second GFP codon, placed the GFP coding sequence in the +1 reading frame relative to the start codon (**Fig. 1a**). Thus, insertions of 3n-1 nucleotides or deletions of 3n-2 nucleotides that occur in the 34 nucleotide leader or the unstructured GFP N-terminus, should restore the expression of a functional GFP. We expanded small founder populations of GFP-negative HCoV-OC43/fsGFP(WT) infectious virions arising from transfection of the cloned viral genome, and after 2 passages (P2) screened viral populations comprising approximately 4x10^7^ plaque forming units (PFU), for mutants exhibiting GFP expression (**Fig. 1b**). Based on the number wells containing GFP+ cells (**Supplementary table 1**), we estimated that the expanded HCoV-OC43/fsGFP populations contained indel mutations that resulted in GFP expression at a frequency of ∼1x10^-7^/PFU.

**Fig.1.**
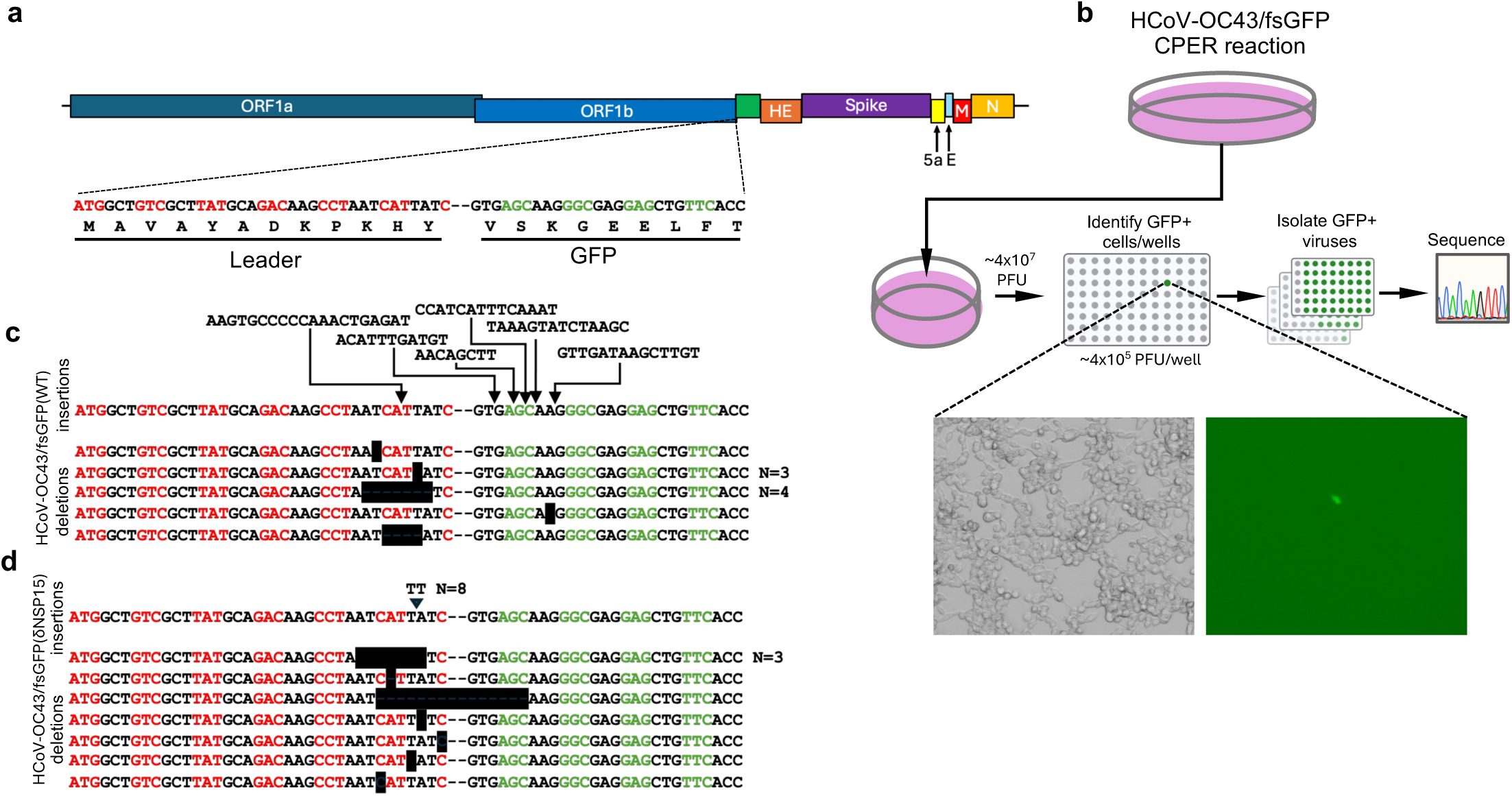
Coronavirus reporter that detects insertion and deletion mutations. (a) Schematic representation of the HCoV-OC43/fsGFP reporter virus with a –2 frameshift in a leader peptide between the start codon and eGFP coding sequence. Sequences of leader peptide (alternating red/black codons) and GFP N-terminus (alternating green/black codons) are shown. (b) Outline of assay used to identify viruses with frameshift mutations. Small founder, GFP-negative virus populations generated by transfection of circular polymerase extension reaction (CPER) product from a molecularly cloned progenitor were expanded (2 passages) and GFP+ viruses isolated by limiting dilution. A magnified view of a well with a GFP+ cell from which an GFP+ viral mutant was obtained is shown. (c,d) Sequences of leader peptide (alternating red/black codons) and GFP N-terminus (alternating green/black codons) in GFP+ viruses isolated following passage of HCoV-OC43/fsGFP(WT) (c) and HCoV-OC43/fsGFP(δNSP15) (d) progenitors. Black boxes indicate deleted sequences, arrows indicate the locations of inserted sequences, insertions or deletions identified in multiple GFP+ virus isolates are marked: “N=” and the number of GFP+ virus isolates with that mutation is indicated.

Using iterative cycles of limiting dilution, we isolated 18 GFP+ viruses from the screened HCoV-OC43/fsGFP(WT) populations (**Fig. 1b**). Sequence analysis of the fsGFP leader and N-terminus in these viruses revealed that all of the isolated GFP-expressing viruses had frameshift mutations in the leader peptide or the unstructured GFP N-terminus that could account for the generation of GFP fluorescence (**Fig. 1c**). Specifically, 10/18 GFP-expressing viruses had 3n-2 nucleotide deletions (7 different species), while the remaining 8/18 (6 different species) had 3n-1, (8 to 20) nucleotide insertions (**Supplementary table 1**).

We repeated the aforementioned experiments using a mutant virus, HCoV-OC43/fsGFP(δNSP15), encoding a presumptively inactivating mutation (H234A) at the active site of NSP15, a conserved ribonuclease expressed by coronaviruses^12^. We isolated a similar number of GFP+ viruses (n=16) from the P2 HCoV-OC43/fsGFP(δNSP15) populations, and GFP expression was again accounted for by frameshift mutations acquired in the leader and GFP N-terminus (**Fig. 1d**). However, in the case of HCoV-OC43/fsGFP(δNSP15), GFP+ mutant derivatives either had deletions or a small insertion (2 nucleotides - that may have been derived from a single P1 founder). None of the GFP+ viruses isolated from the HCoV-OC43/fsGFP(δNSP15) parent had inserts >2 nucleotides (**Supplementary table 1**).

### Deep sequencing of HCoV-OC43 genomes reveals NSP15-dependent insertion mutations

To quantify insertion mutations and the effect of NSP15 on their generation, we next produced HCoV-OC43/fsGFP (WT) and mutant HCoV-OC43/fsGFP (δNSP15) virus stocks, harvested after 2 passages following transfection of cloned viral DNA. We subjected RNA collected from purified virions (8.2 x10^8^ PFU for WT and 2.4x10^8^ PFU for δNSP15) to ultra-deep short-read sequencing. This approach generated 6.4x10^8^ (WT) and 4.2x10^8^ (δNSP15) virus-derived sequencing reads, resulting in a sequencing depth of 10^6^x to 10^7^x across nearly the entire viral genome (**Extended Data Fig. 1a**). We analyzed the reads using an algorithm configured to detect insertions, resulting in the detection of 21894 and 2925 unique insertions of 2 nucleotides or larger into the WT and δNSP15 viral genomes, respectively (**Supplementary dataset 1**). These values correspond to 33.9 and 6.93 event counts per million reads (cpm), or ∼6.8x10^-3^ and 1.4x10^-3^ inserts per viral genome. In addition to the discrepancy in the number of inserts detected for WT and δNSP15 viruses, analysis of the insert size distribution revealed a striking difference. Most inserts in the HCoV-OC43/fsGFP(WT) were in the range 5 to 30 nucleotides (median 12 nucleotides) while those in HCoV-OC43/fsGFP(δNSP15) were 2-3 nucleotides (median 3 nucleotides, **Fig. 2a**). Thus, consistent with the results using HCoV-OC43/fsGFP indicator virus (**Fig. 1**) sequencing of viral populations demonstrated that NSP15 greatly increases the number of insertion mutations of >2 nucleotides into the HCoV-OC43 genome.

**Fig. 2.**
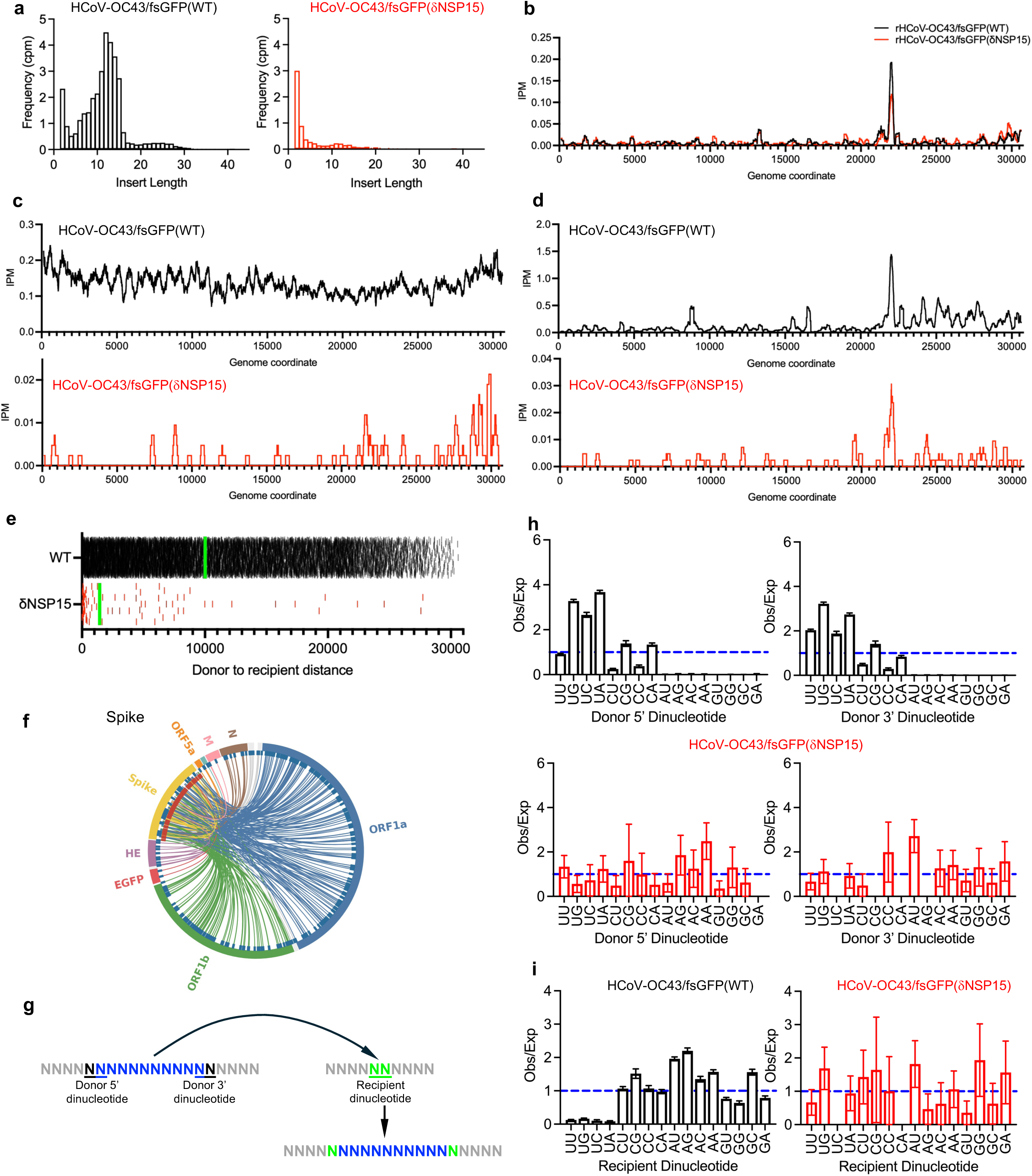
Ultra deep sequencing of HCoV-OC43 virions reveals NSP15-dependent insertion mutations. (a) Frequency distribution of insert lengths (in nucleotides) for all inserts of 2 nucleotides or longer in sequencing reads, given as counts per million sequencing reads (cpm), from RNAseq analysis of purified HCoV-OC43/fsGFP virions, WT (left, black) and δNSP15 (right, red). (b) Frequency of tandem (side-by-side) duplications in a sliding window of 250 nucleotides across the WT (black) and δNSP15 (red) HCoV-OC43/fsGFP genomes, for inserts of probable viral origin, given as inserts per 250 nucleotides per million reads (IPM). (c, d) Frequency with which sequences within a sliding window of 250 nucleotide across the WT (upper, black) and δNSP15 (lower, red) HCoV-OC43/fsGFP genomes, serve as donor (c) and recipient (d) sites for distal inserts of probable viral origin, given as inserts per 250 nucleotides per million reads (IPM). (e) Distances (in nucleotides) between donor and recipient sites for distal inserts of probable viral origin for WT (black) and δNSP15 (red) HCoV-OC43/fsGFP sequences in purified virion RNA. Each point represents a different insertion event, green lines = median distance between donor and recipient sites. (f) Circos plot, indicating locations of 320 donor sequences (blue ticks, outer circle) and 160 recipient sites (red ticks, inner circle) for chimeric inserts with two viral insert donors into each recipient site in the spike gene of HCoV-OC43/fsGFP(WT). (g) Schematic indicating 5’ and 3’ donor dinucleotides (blue/black) and recipient dinucleotides (green), that were analyzed for compositional bias. (h) Observed/expected (Obs/Exp) frequency of the occurrence of dinucleotides at the 5’ and 3’ ends of inserts in the context of the donor sites for distal inserts of probable viral origin for WT (black, upper) and δNSP15 (red, lower) HCoV-OC43/fsGFP sequences in purified virion RNA. Inserts with ambiguous insertion sites were not included. (i) Observed/expected (Obs/Exp) frequency of the occurrence of dinucleotides at unambiguous recipient sites for distal inserts of probable viral origin for WT (black, left) and δNSP15 (red, right) HCoV-OC43/fsGFP sequences in purified virion RNA.

Analysis of *k*-mers derived from the positive and negative sense HCoV-OC43 RNA indicated that random 8mer, 9mer and 10mer oligonucleotides have a ∼48%, ∼19% and ∼5.4% probability of aligning to HCoV-OC43/fsGFP sequences by chance (**Supplementary table 2**). Thus, to assess the origin of the viral inserts with minimal ambiguity, we confined our initial analysis to inserts of 11 nucleotides or larger (11+ nucleotides) that have a <5% probability of occurence in the viral genome by chance. Of the 11+ nucleotide inserts, 12063/13816 (87%) in the WT genome and 508/655 (77%) in the δNSP15 genome, had a perfect or near perfect match to sequences in the viral genome and were considered to be of likely viral origin (see methods). The majority of these presumptively viral genome-derived inserts (90% WT, 99% δNSP15) corresponded to plus-strand viral sequences (**Supplementary table 3**).

This set of 11+ nucleotide inserts with a definable viral origin and destination site in the viral genome fell into three main classes (**Extended Data Fig. 1b**). A minority class (5.2%) consisted of tandem, or “side-by-side” duplications, in which an insert sequence matched that of the viral genome immediately proximal to the insert. The location (**Fig. 2b**), frequency (0.98 vs 0.99 cpm) and size (median = 17 vs 16 nucleotides, **Extended Data Fig. 1c**), of these inserts was similar for WT and δNSP15 viruses.

The majority of inserts fell into a second class, that could be mapped to the viral genome and had an apparent viral origin site that was distal to destination site (**Extended Data Fig. 1b**). These distal inserts were far more numerous (84x) in the WT (16.7 cpm) than the δNSP15 (0.2 cpm) sequencing datasets (**Fig. 2c,d**). The median size of these inserts was 13 (WT) and 15 (δNSP15) nucleotides (**Extended Data Fig. 1d**). The donor sequences of the distal inserts corresponded to oligonucleotides that were apparently randomly distributed across the HCoV-OC43/fsGFP(WT) genome (**Fig. 2c**), while the recipient sites of insertion were concentrated toward the 3’ end of the genome and were particularly frequent in the embedded GFP reporter gene (**Fig. 2d**). In the case of HCoV-OC43/fsGFP(WT), the distribution of distances between insert donor and recipient sites (median= ∼10kB) was similar to that expected for two randomly selected locations on the ∼30kB viral genome (**Fig. 2e Extended Data Fig. 1e**). In contrast, for HCoV-OC43/fsGFP(δNSP15), the location of insert donor and recipient sites was strongly biased so that the two sites were close to each other (**Fig. 2e**).

Overall, these data suggest at least two distinct modes of insert generation in HCoV-OC43 sequence datasets. The dominant mechanism, in which the insert donor and recipient sites are apparently unlinked, was strongly dependent the presence of NSP15 during viral replication.

A third insert class was similar to the second, but drew on two different, distal viral donor sites to generate chimeric inserts (**Extended Data Fig. 1b**). Because minimally ambiguous assignment of inserts to this class required the identification of two different 11+ nucleotide sequences within a single insert and are thus, by definition, >22nt long, our catalogue represents an underestimate of the total numbers of inserts in this class. Nevertheless, we identified 467 chimeric inserts (3.7% of all 11+ nucleotide viral origin inserts) (**Fig. 2f**, **Extended Data Fig. 2**). These chimeric inserts were found in all viral genes, but exclusively in the HCoV-OC43/fsGFP(WT) dataset and were not detected in HCoV-OC43/fsGFP(δNSP15).

### Insert sequences suggest NSP15-dependent genesis

At least two hypotheses could explain the requirement for NSP15, a nuclease, in the generation of distal virus-derived inserts into the viral genome. Specifically, NSP15 could be required (i) for the generation of the inserted sequences, or (ii) for the generation of cleavage sites at which the inserts are incorporated into the viral genome. We inspected dinucleotides at the −1 and +1 positions at the 5’ and 3’ boundaries of insert donor sites as well as −1/+1 dinucleotides at recipient sites (**Fig. 2g**). Note however, that the identity of these dinucleotides can be ambiguous in cases where the same base is present at the end of the insert and at the recipient site (**Extended Data Fig. 3**), and such ambiguous instances were excluded from the analyses below. These analyses revealed clear biases. Specifically, both the 5’ and 3’ ends of the inserts in HCoV-OC43/fsGFP(WT) were derived most frequently from sequences that had UN dinucleotides at the −1/+1 positions at the 5’ and 3’ ends of the insert donor sequence (**Fig. 2h**). Conversely, AN and GN dinucleotides were strongly disfavored at these dinucleotide positions. These dinucleotide biases are similar to the reported substrate preference of NSP15 nucleases encoded by SARS-CoV-2 and MHV^13,14^. No such dinucleotide biases were observed for the smaller number of inserts found in HCoV-OC43/fsGFP (δNSP15) (**Fig. 2h**). Inspection of sequences flanking the recipient sites, where inserts were acquired in the viral genome, did not reveal any biases that would suggest a role for NSP15 in cutting the genome to generate the recipient site. Rather, UN dinucleotides that constitute preferred NSP15 cleavage sites were underrepresented at recipient sites for the HCoV-OC43/fsGFP(WT) insert dataset (**Fig. 2i**). Again, no such biases were observed for the HCoV-OC43/fsGFP(δNSP15) inserts.

Based on these analyses, we reexamined the GFP+ viruses isolated from the HCoV-OC43/fsGFP(WT) biological assay (**Fig. 1c**). Of the 6 insertions of >3nt found therein, 5 could be completely or partially mapped to a single location in the viral genome uniquely, while the remaining 1 insert could be mapped to 6 locations. (**Fig. 1a, c**, **Supplementary Table 1**) Overall, these data indicate that most distal inserts in the HCoV-OC43/fsGFP(WT) viral genome are derived from the products of RNA cleavage by NSP15.

### NSP15 as a source of genetic innovation in SARS-CoV-2

To determine whether NSP15 was similarly required for the generation of insertion mutations in SARS-CoV-2, we analyzed public domain datasets derived from deep sequencing WT and NSP15-mutant SARS-CoV-2 virion RNA^19^. The methods for generating this published dataset were different to those used herein for HCoV-OC43/fsGFP, and the datasets had less depth (108 million and 98 million mapped reads for SARS-CoV-2(WT) and SARS-CoV-2(δNSP15), respectively). Nevertheless there were 1705 and 1393 unique insertions of 2 nucleotides or larger into the SARS-CoV-2(WT) and SARS-CoV-2 (δNSP15) viral genomes respectively (**Supplementary Table 3, Supplementary dataset 2**), corresponding to 15.8 and 14.2 event counts per million reads (cpm), or 3.2x10^-3^ and 2.82x10^-3^ inserts per viral genome. Notably, SARS-CoV-2(WT) had a greater proportion of inserts in the range 5 to 30 nucleotides while those in SARS-CoV-2(δNSP15) were dominated by 2-3 nucleotide inserts (**Fig. 3a**). These differences were particularly evident when analysis was confined to inserts of 11+ nucleotides whose origins could be mapped to the viral genome. (**Extended Data Fig. 4a**).

**Figure 3.**
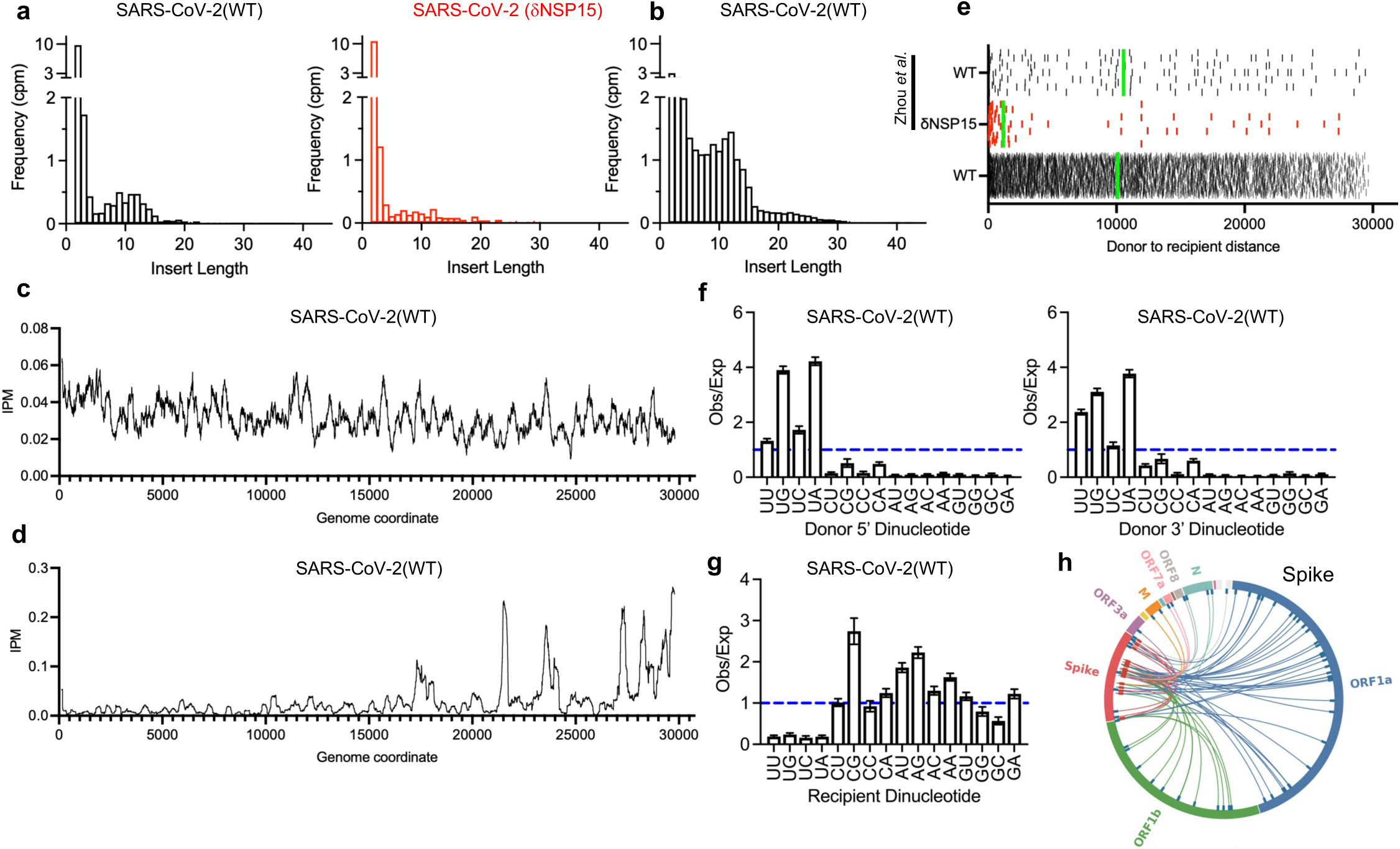
Deep sequencing of SARS-CoV-2 (WT) and (δNSP15). (a,b) Frequency distribution of insert lengths (in nucleotides) for all inserts of 2 nucleotides or longer in sequencing reads, given as counts per million sequencing reads (cpm) from published sequencing datasets from SARS-CoV-2(WT) (left, black) and δNSP15 (right, red) (a) or from a RNAseq datasets from SARS-CoV-2(WT) virions with 991,880,264 mapped reads analyzed using the same methods as in Fig. 2 (b). (c, d) Frequency with which sequences within a sliding window of 250 nucleotide across the SARS-CoV-2 genome, serve as donor sites (c) or recipient sites (d) for distal inserts of probable viral origin, given as inserts per 250 nucleotides per million reads (IPM). (e) Distances (in nucleotides) between donor and recipient sites for distal inserts of probable viral origin for WT (black) and δNSP15 (red) SARS-CoV-2 sequences for published datasets (Zhou et.al.) and purified virion sequencing datasets generated herein. Each tick represents a different insertion event, green lines = median distance. (f, g) Observed/expected (Obs/Exp) frequency of the occurrence of dinucleotides at the 5’ and 3’ ends of inserts in the context of unambiguous insert donor sites (f) and recipient sites (g) for distal inserts of probable viral origin in SARS-CoV-2 sequences from purified virion RNA generated herein. (h) Circos plot, indicating locations of 66 donor sequences (blue ticks, outer circle) and 33 recipient sites (red ticks, inner circle) for chimeric inserts with two viral insert donors into each recipient site in the spike gene of SARS-CoV-2.

Because all published SARS-CoV-2 sequencing datasets, including the aforementioned, have considerably less depth than is optimal for analysis of rare events such as insertional mutation, we generated a deeper sequencing dataset for SARS-CoV-2(WT) using methods similar to those used above for HCoV-OC43/fsGFP (**Fig. 2**). Specifically, a SARS-CoV-2 virus stock was generated from cloned SARS-CoV-2 DNA by CPER transfection and expansion over two passages. Then, RNA was extracted from purified viral particles, corresponding to 3.6x10^7^ PFU. Analysis of 9.9x10^8^ reads from this viral population gave sequence coverage that was similar to the HCoV-OC43/fsGFP analysis (10^6^x to 10^7^x at each genome position, **Extended Data Fig. 4b**). This dataset yielded 21406 total inserts of >2 nuclotides (corresponding to an insert frequency of 4.3x10^-3^ per viral genome. The median insert length in SARS-CoV-2 (8 nucleotides) was shorter than in HCoV-OC43/fsGFP (**Fig. 3b**) but there were, nevertheless, 7,371 inserts of 11+ nucleotides, of which 6141 (83%) could be mapped to the SARS-CoV-2 genome with minimal ambiguity (**Supplementary Table 3, Supplementary dataset 3**). Of the 11+ nucleotide inserts of probable viral origin, 3772 constituted distal insertions while 2183 represented tandem duplications (**Extended Data Fig. 4c**). Donor sites for distal insertions were distributed throughout the SARS-CoV-2 genome (**Fig. 3c**), while distal insert recipient sites were concentrated in regions that were more often in the 3’ portion of the viral genome (**Fig. 3d**) and sometimes overlapped with regions that were enriched in tandem duplications (**Extended Data Fig. 4d**). The distances between donor and recipient sites in both ultra deep and published SARS-CoV-2(WT) datasets was as expected for randomly selected pairs of points on the SARS-CoV-2 genome (**Fig. 3e, Extended Data Fig. 1e**), while donor and recipient sites were more closely linked for SARS-CoV-2(δNSP15). In each of these properties, the distal SARS-CoV-2 inserts resembled the HCoV-OC43/fsGFP inserts in terms of the effect that NSP15 exerted on their characteristics. Additionally, the 5’ and 3’ ends of the distal inserts in both the larger SARS-CoV-2(WT) dataset and the published SARS-CoV-2(WT) dataset were primarily derived from donor sequences that had UN dinucleotides at the −1/+1 positions at both ends, while UN dinucleotides were strongly disfavored at insert recipient sites (**Fig. 3f,g Extended Data Fig. 4e**). The smaller number of distal inserts in SARS-CoV-2(δNSP15) did not exhibit these characteristics. Chimeric inserts derived from two different viral donor sites were also a feature of SARS-CoV-2(WT), with 114 distal inserts being clearly chimeric (**Fig. 3h, Extended Data Fig. 5**). Again, such characteristics were similar to the inserts found in HCoV-OC43/fsGFP(WT), and these data indicate that NSP15-driven insertional mutagenesis, while marginally less active in SARS-CoV-2 as compared to HCoV-OC43, represents a major source of genetic innovation in both of these coronaviruses.

### Insertion mutations in SARS-CoV-2 RNA during human and mouse infection

We next examined ultra-deep sequencing libraries generated from PCR amplicons, rather than libraries generated directly from purified virion RNA, noting that PCR amplification can generate occasional sequence rearrangements that might conflate detection of insertion mutations^20,21^. We generated a 162 nucleotide PCR amplicon (116 nucleotides after primer removal; covering ∼0.4% of the viral genome) spanning the S1/S2 junction, using a complex source consisting of 7.8x10^9^ RNA genomes from purified SARS-CoV-2 virions. As a control, we also generated the same amplicon using the most dilute virion RNA sample that yielded PCR product (∼2 RNA molecules). PCR generated artifacts should appear in both amplicons, while insertion mutations resulting from viral replication should be present in the first but absent (or nearly absent) from the second amplicon. We found tandem duplications of 11+ nucleotides (n= 39) among 3.45 x10^8^ reads and additional, rare, apparently intra-amplicon micro-rearrangements (n=2) similar to those previously reported to occur during PCR ^20^ ^21^ in the 2-molecule amplicon. These artifacts could be filtered by applying our previous criteria (by considering insertions of 11+ nucleotides of probable viral origin, **Supplemental table 2**), and applying the addional constraint that inserts originate from viral sequences outside the amplicon. By these criteria, we found 876 inserts in the 3.8x10^8^ reads of the 116 nucleotide amplicon derived from high complexity virion RNA (**Fig. 4a**, **Supplementary dataset 4**), and 0 such inserts in the control amplicon, derived from ∼2 RNA template molecules.

**Figure 4.**
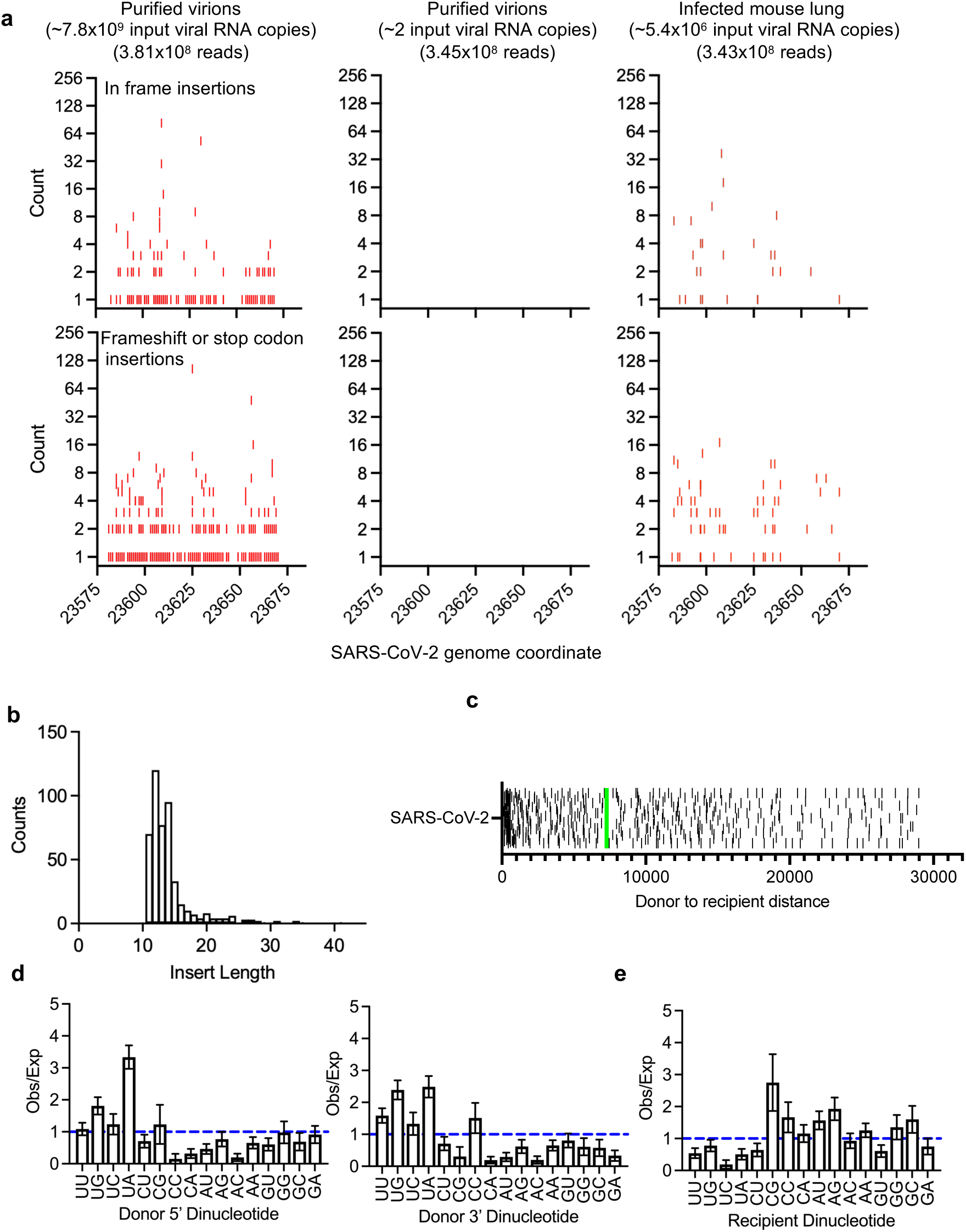
Analysis of SARS-CoV-2 insertional mutants in various contexts. (a) Charts, with the SARS-CoV-2 genome position represented on the X-axis, showing the number of read counts (Y axis) in which individual insertional mutants in a 116 nucleotide amplicon were found. Amplicons were generated using samples of purified virion RNA or infected mouse lung RNA containing the indicted number of viral RNA copies. Each individual red tick represents an insertion event that generates in frame insertions (upper) or protein truncations (lower). (b) Frequency distribution of insert lengths (in nucleotides) for distal inserts of probable viral origin of 11 nucleotides or longer in sequencing reads from a published sequencing dataset (PRJNA613958) from SARS-CoV-2 infected humans, given as raw counts. (c) Distances (in nucleotides) between donor and recipient sites for distal inserts of probable viral origin from published sequencing dataset (PRJNA613958) from SARS-CoV-2 infected humans. Each tick represents a different insertion event, green lines = median distance. (d, e) Observed/expected (Obs/Exp) frequency of the occurrence of dinucleotides at the 5’ and 3’ ends of inserts in the context of unambiguous insert 5’ and 3’ donor sites (d) and recipient sites (e) for distal inserts of probable viral origin from published sequencing dataset (PRJNA613958) from SARS-CoV-2 infected humans.

We applied the same PCR amplicon approach to RNA harvested from SARS-CoV-2 infected K18/hACE2 mouse lungs at 3 days after intranasal infection with 20,000 PFU of SARS-CoV-2. This viral population was obviously smaller than that sampled as the purified virions. Nevertheless, from an estimated (2.7x10^6^) input viral RNA molecules from infected mouse lung RNA, we found 95 inserts of 11+ nucleotides and probable viral origin (**Fig. 4a, Supplementary dataset 4**).

Next, we analyzed two sets of SARS-CoV-2 sequences obtained from two clinical datasets in public databases (PRJNA613958, PRJNA805055, **Supplementary Table 4, Fig. 4 b-e**, **Extended Data Fig. 6a-d**). These sequences were generated after PCR amplification from clinical samples using amplicons covering nearly the entire viral genome. Again, we confined our analysis to distal insertion mutations of 11+ nucleotides in which the insert was not derived from sequences in the same amplicon. Analysis of 488 and 301 distal inserts (**Supplementary dataset 5**) found in these two clinical datasets revealed a size distribution and donor-to-recipient distance which resembled that accompanying NSP15 mediated processes described in purified HCoV-OC43 and SARS-CoV-2 virions (**Fig. 4b,c**, **Extended Data Fig. 6a,b**). Analysis of donor and recipient dinucleotides also revealed an overrepresentation of UN dinucleotides, and underrepresentation of AN and GN at insert donor 5’ and 3’ dinucleotide ends (**Fig. 4 d**, **Extended Data Fig. 6c**). Conversely, UN dinucleotides were underrepresented at recipient sites. (**Fig. 4e**, **Extended Data Fig. 6d**). These insert characteristics again resembled, but were less extreme than, those associated with distal inserts in the purified HCoV-OC43 and SARS-CoV-2 virions, perhaps reflecting an additional superimposed mechanism of insert generation during human infection, or contamination of the clinical sequence dataset by PCR-induced rearrangements.

### Frequent acquisition of potential furin cleavage sites in coronaviruses

Herein, we documented thousands of insertion mutations in coronavirus genomes acquired during normal virus replication. While the majority of such quasi-randomly generated insertion mutations are expected to be either lethal, or carry a fitness penalty, a small fraction may be beneficial and enhance viral replication. We first quantified the insertion mutations in open reading frames that were in frame versus those that caused frameshifts or truncations. For HCoV-OC43, 3794/13183 (28%) of inserts in open reading frames were in frame (**Extended Data Fig. 7**), while for SARS-CoV-2 1501/4952 (30%, dataset generated herein, **Fig. 5a**) and 63/216 (29%, published dataset^19^, **Extended Data Fig. 8**) were in frame. The inserts encoding frameshifts or truncations were not obviously enriched in non-essential genes, and the degree of their preponderance is close to that expected for random sequence inserts. These data thus indicate that minimal selection pressure has acted upon most inserts, and suggest that the genomes with inserts are either newly generated, or carried as low frequency passengers in the viral population by intact complementing viral species.

**Figure 5.**
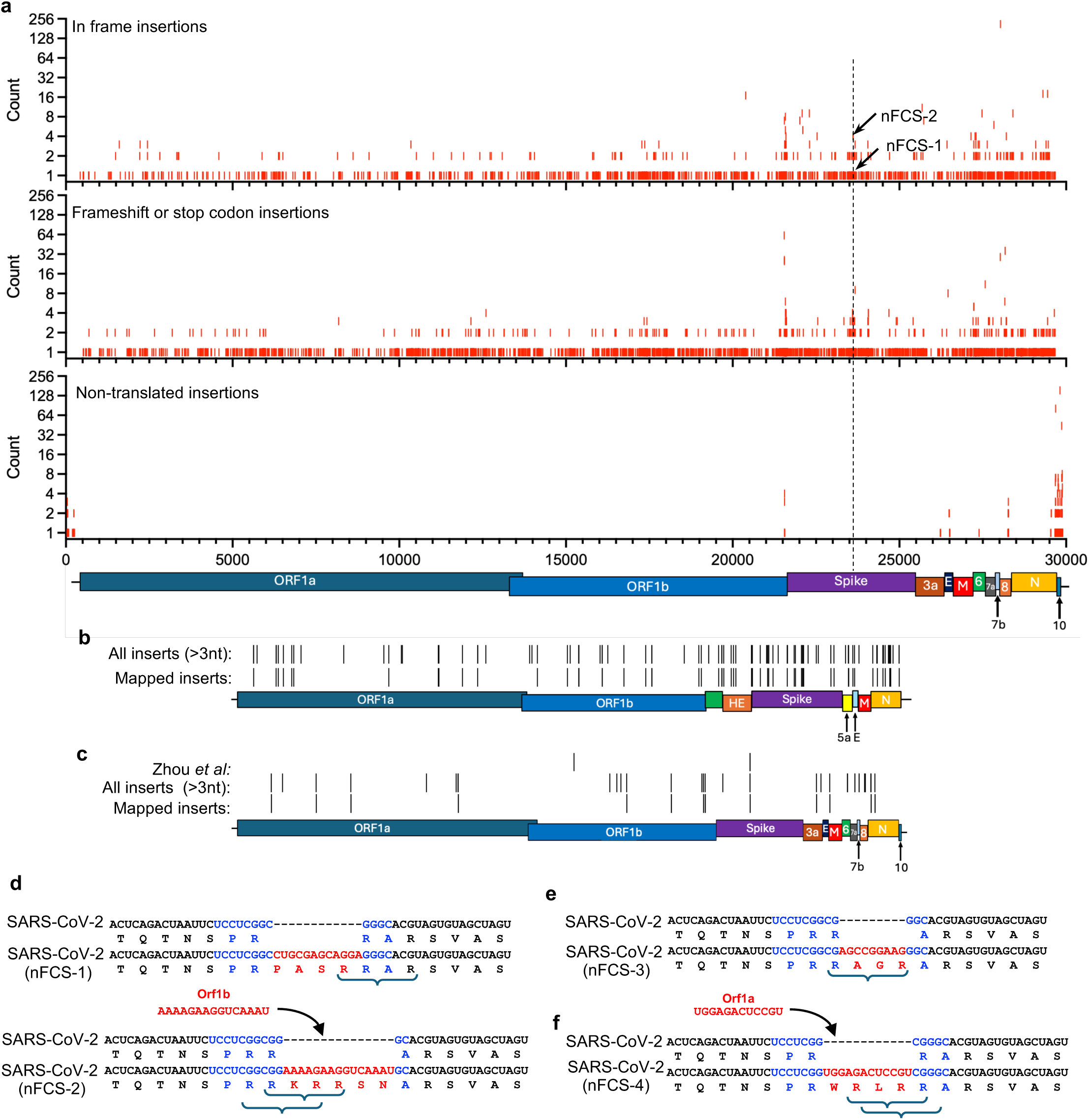
Coding potential of insertional mutations and recurrent acquisition of furin cleavage sites during coronavirus replication. (a) Charts, with the SARS-CoV-2 genome position represented on the X-axis showing the number of reads on the Y axis (out 991,880,264 total mapped reads) in which individual insertional mutants in the sequenced virion population that generate in frame insertions (upper), protein truncations (middle), or occurred in untranslated regions (lower). Each individual red tick represents an insertion event. Datapoints corresponding to the appearance of two new furin cleavage sites (nFCS-1 and −2), at the site of the existing SARS-CoV-2 spike furin cleavage (vertical dashed line) site are highlighted. (b, c) Schematic representation of HCoV-OC43/fsGFP genome (b) and SARS-CoV-2 genome (c), with indicated positions of insertion mutations that result in the acquisition of potential furin cleavage sites in the sequenced viral populations or in a published SARS-CoV-2 dataset (Zhou et al). “All inserts >3 nucleotides” included viral derived and ambiguous origin inserts, “mapped inserts” represent inserts of probable viral origin. (d-f) Acquisition of novel furin cleavage sites (nFCS-1, −2 −3 and −4), at the site of the existing SARS-CoV-2 furin cleavage site, during the course of SARS-CoV-2 replication. The pre-existing SARS-CoV-2 FCS insertion is shown in blue, sites acquired during replication of molecularly cloned virus are shown in red. The nFCS-1, −2 (d) were found in the SARS-CoV-2 population sequenced herein, nFCS-3 (e) was found in a published SARS-CoV-2 dataset (Zhou et al) while nFCS-4 (f) was found in the purified virion PCR amplicon dataset. Potential furin cleavage sites (RXXR motifs) are bracketed.

An example of genetic innovation arising from insertion mutations that may in some circumstances confer increased fitness, is the acquisition of new potential furin cleavage sites (FCS). Indeed, an insertion mutation at the S1/S2 junction in the SARS-CoV-2 spike protein, relative to its sarbecovirus ancestors, that generates an FCS and enhances viral transmissibility may, in part, be responsible for the SARS-CoV-2 pandemic. The minimal requirement for a potential FCS is “RXXR” and because arginine is specified by 6 out of the 64 codons in the universal genetic code, roughly 0.9% of random 12 nucleotide sequences are expected to encode a potential FCS. Thus, we found numerous examples of potential FCS generation through insertional mutagenesis in our sequencing datasets. Specifically, in purified HCoV-OC43 and SARS-CoV-2 virions, inserts acquired during virus replication generated 108 and 41 new species with potential FCS, respectively (**Fig. 5b, c, Supplementary datasets 1, 3**). These FCS were apparently randomly distributed among the viral proteins and most are likely functionally irrelevant. However, we identified numerous examples of FCS acquisition through insertional mutagenesis, precisely at the location of the existing FCS in SARS-CoV-2 (**Fig. 5d-f Extended Data Fig. 9**). Two of these new (n) FCS, (nFCS-1 and −2) were found in the sequencing dataset generated herein. A third, nFCS-3, was found in the published dataset^19^. Interestingly, nFCS-3 increased in frequency during viral passage (0/28 million reads in P1 virus stock vs 774/80 million reads at a later passage)^19^ suggesting a replication advantage. One of the mutants in our dataset (nFCS-2) encoded a high basic peptide RRKRR, with 2 potential overlapping FCS, that was generated by insertion of 15 nucleotide sequence from Orf1b, while nFCS-1 and nFCS-3 were of ambiguous, possibly chimeric origin (**Fig. 5d, e**). Analysis of PCR amplicon datasets in which sequences spanning S1/S2 junction were selectively amplified from purified SARS-CoV-2 viron RNA (**Fig. 4a**), revealed that 15/876 (1.7%) of the 11+ nucleotide inserts of probable viral origin found in this 116 nucleotide region generated new or alternative FCS, precisely at the location occupied by the exisiting SARS-CoV-2 FCS (**Fig. 5f, Extended Data Fig. 9a**). Simililarly 2/95 (2.1%) of the probable viral inserts in this region sequenced from infected mouse lung RNA generated replacement nFCS sites at the site of the existing FCS (**Extended Data Fig. 9b**). In considering the origins of the exisiting FCS in SARS-CoV-2, models that could account for its generation (**Extended Data Fig. 10**) include chimeric inserts of viral origin, of the type observed in our datasets (**Fig. 2f**, **Fig. 3h**) which give intermediates that differ by a single nucleotide from the prototypic SARS-CoV-2 sequence. We also note the presence of sequences in human RNA that perfectly match to the 12 nucleotide SARS-CoV-2 insert^22^, and the fact that a fraction of the inserts found in our purified SARS-CoV-2 virion datasets (0.2%) appear derived from host sequences (**Supplementary table 3**).

## Discussion

Our data suggest a model in which the digestion of coronavirus RNA by NSP15 generates a pool of oligonucleotides that are subsequently acquired as insertions into the viral genome during its replication. While it is formally possible that such oligonucleotides might be directly ligated into the viral genome, we favor a model in which diffusible RNA oligonucleotides invade the elongating polymerase complex during genome replication, serving as temporary templates through elongating strand-transfer. Most virus-derived inserts had a polarity matching that of the viral genome (+ sense in virion RNA), and since + strands are more abundant than – strands in infected cells^23,24^ this finding suggests that inserts are acquired by polymerase transfer from the + strand genomic RNA template to + strand oligonucleotide templates during – strand synthesis. The clear enrichment of favored NSP15 target dinucleotides at insert donor sites obviously suggests that NSP15 is directly responsible for generating the RNA oligonucleotides that give rise to inserts. The paucity of favored NSP15 dinuclotide targets at recipient sites is more difficult to explain, but hints at the possibility that NSP15 might preferentially cleave inserts that become incorporated immediately 3’ to U nucleotides.

In addition to the clear NSP15 driven insertional mutational process, our datasets contain tandem duplications in which sequence inserts into virion RNA that match the neighboring portion of the viral genome are found. The appearance of such inserts in both WT and δNSP15 datasets renders their origin ambiguous. Moreover, our finding that tandem duplications are found in a PCR amplicon dataset derived from a low complexity sample (∼2 RNA molecules) indicates that some tandemly duplicated inserts can be generated as artifacts during PCR amplification or sequencing library generation. Conversely, the absence of distal inserts in amplicons prepared from a low complexity RNA sample, coupled with the large increase in their number when NSP15 was present during viral replication, as well as their independent discovery as fixed mutations in viruses isolated through the biological GFP frameshifting assay, clearly indicates the natural provenance of such inserts.

Estimates of the frequency with which insertional mutants occur in a viral population are predicated on knowledge of the size of the viral population analyzed. Based on the library preparation procedure deployed herein, the sequencing depth deployed (10^8^-10^9^ reads of ∼150 nucleotides) was quite well matched to the number of viral RNA molecules sampled during sequencing library preparation (∼3x10^7^ copies of a ∼30,000 RNA genome), enabling an estimate of insertion mutant frequency that we determine to be in excess of 10^-3^ per genome. Conversely, clinical sequence datasets derived from PCR amplicon sequencing are of unknown and likely lower complexity, with selective pressures potentially purging largely deleterious insertional mutations. Thus, estimates of insertional mutant frequency cannot be made from clinical datasets. Nevertheless, our analysis of sequences from published clinical sample datasets indicates the occurrence of NSP15-driven mutagenic processes, albeit possibly superimposed with other mutational processes or amplification/sequencing artifacts.

The acquisition of an FCS in the spike protein of SARS-CoV-2 that is absent in its closest relatives that circulate in *rhinolophus* bats has been the subject of much controversy. Lack of knowledge about the frequency and mechanism by which insertion mutations occur in RNA viruses has led some commentators to cite its presence as evidence of human manipulation^25^. However, because RNA virus replication often generates extremely large populations, the occurrence of events that might intuitively seem unlikely, or go undetected in hundreds of viral genome sequences, is in fact, commonplace during the normal course of viral propagation in nature. The acquisition of FCS in the spike protein of SARS-CoV-2 is an example of such an event. Indeed, in a single modestly sized SARS-CoV-2 population, we found numerous insertions that generate new FCS-like sequences. Therefore, the emergence of coronaviruses with FCS at the S1/S2 boundary is largely a function of host environment-dependent selective pressures, and is not limited by the frequency with which they are generated through natural mutational processes.

## Methods

### Cells and virus titration

293T-ACE2^26^, 293T (ATCC; #CRL-3216), and VeroE6 (gift from Ralph Baric) cells were cultured in Dulbecco’s Modified Eagle Medium (DMEM) supplemented with 10% fetal calf serum (FCS; Sigma; #F0926) and 10 µg/mL gentamicin (Thermo-fisher; #15750078) at 37°C and 5% CO_2_. Cells were checked periodically for mycoplasma and retrovirus contamination by Hoeschst staining and reverse transcriptase assays. All coronavirus infections were done at 34°C and in the presence of 5% CO_2_. Virus titers were quantified using standard plaque assays. HCoV-OC43 titers were measured on 293T cells and SARS-CoV-2 titers were measured on VeroE6 cells. Plates (24 well) containing 200,000 cells were inoculated with 10x serial dilutions of virus in 200 µL serum free DMEM for 90 minutes. Inoculum was aspirated and 0.5mL DMEM (with 2% methylcelluose, 2% FCS, and 10 µg/mL gentamicin) was added per well. After 6 days, the overlay was removed, wells were washed with PBS twice, cells were fixed with 4% paraformaldehyde for 15 minutes and washed with PBS. The GFP fluorescence in SARS-CoV-2/dORF7/GFP infected cells was used to quantify titers after imaging on an EVOS m7000 microscope (Thermo-Fisher). For HCoV-OC43 infected cells, the monolayer was incubated with PBS containing 10% goat serum (Sigma; #G9023-5ML) and 0.1% Triton-X100 for 30 minutes. Cells were then incubated with mouse anti-HCoV-OC43 N (Sigma #MAB9013) diluted 1:1000 in antibody dilution buffer (10% goat serum and 0.1% Tween-20 in PBS), washed three times with PBS, and incubated with an anti-mouse secondary antibody conjugated to a 488 fluorophore (Thermo-Fisher #A11029), and washed three times in PBS. Plaques were imaged on an EVOS m7000 microscope.

### Recombinant coronavirus genomes

The HCoV-OC43 genetics system was based on the previously published circular polymerase extension reaction (CPER) methodology developed for for SARS-CoV-2^27^. Infectious HCoV-OC43 virus (ATCC, #VR-1558) was obtained from Zeptometrix (#0810024CF) and viral RNA was column purified (Macherey-Nagel, #740956). Then, cDNA was prepared using SuperScript III Reverse Transcriptase (Thermo-Fisher, #18080044) which served as a template to amplify the viral genome in 8 fragments using PrimeStar GXL polymerase (Takara, #R050B) and gene specific primers (**Supplementary Table 5**) which were inserted into the CSIN plasmid. A further plasmid containing a linker fragment with overlaps to the 5’ and 3’ end of the HCoV-OC43 genome and containing a cytomegalovirus immediate early gene promoter, bovine growth hormone poly(A) signal, and hepatitis delta virus ribozyme cleavage sequence that served to circularize the recombinant DNA and drive gene expression was also generated. The SARS-CoV-2 CPER system was constructed as previously described^27^ using viral gene fragments synthesized by Geneart.

The HCoV-OC43/GFP virus was created by inserting an GFP gene between the first 13 and last 27 amino acids of the NS2 coding sequence in the CSIN-OC43-Frag6 plasmid to create CSIN-OC43-Frag6/ΔNS2GFP. The HCoV-OC43/fsGFP reporter virus was created by engineering a two base pair deletion in the open reading frame upstream of the GFP coding sequence in CSIN-OC43-Frag6/ΔNS2GFP plasmid. The HCoV-OC43/fsGFP(δNSP15) H234A mutant virus was created using site directed mutagenesis. For SARS-CoV-2/δORF7/GFP, the entire coding sequence of ORF7a was replaced by the coding sequence of eGFP. Information on oligos and restriction enzymes used for cloning is provided in **Supplementary table 5**. All cloning was performed with NEBuilder HiFi DNA Assembly Master Mix (NEB, # E2621).

### Generation of coronaviruses from recombinant DNA

PCR fragments for HCoV-OC43 and SARS-CoV2 were generated by using Primestar GXL DNA polymerase and primers that introduced 40-50 nucleotide homologies between each adjacent fragment in the virus genome (**Supplementary table 5**).^27^ PCR products were extracted from agarose gels, and column purified (Macherey-Nagel, # 740609.250). For the CPER reaction, 50 μL PCR reactions were assembled with 10uL 5X GXL buffer, 4μL dNTP mix, 1 μL of enzyme and 100 ng of each of the fragments for the respective virus.

Volume was adjusted to 50 μL using nuclease-free water. Conditions used were as follows: Initial denaturation at 98°C/5 minutes followed by 35 cycles of 98°C/15s, 55°C/15s, 68°C/25 minutes and a final extension of 68°C for 30 minutes followed by cooling at 4°C. For virus reconstitution, 25 μL of CPER mix was incubated with 8 μL PEI in serum-free DMEM for 10-15 minutes and directly transfected on 293T (HCoV-OC43) or 293T-ACE2 cells (SARS-CoV2). For SARS-CoV-2, transfected 293T-ACE2 cells were co-cultured with VeroE6 cells at 3-4 days post transfection. Virus supernatant was harvested upon observation of cytopathic effect,.

### HCoV-OC43/fsGFP revertant assay

Passage one (P1) stocks of HCoV-OC43/fsGFP and HCoV-OC43/fsGFP(δNSP15) were recovered from 293T cells following transfection with the CPER reaction products. Then, 293T cells (400,000 in one well of a 12 well plate) were infected with HCoV-OC43/fsGFP or HCoV-OC43/fsGFP(δNSP15) P1 viruses at a multiplicity of 3 plaque forming units (PFU) per cell. At 24h after infection, supernatant was harvested, filtered (0.22µm) and 1mL containing ∼4x10^7^ PFU was distributed over wells of a 96 well plate, each well containing 20,000 293T cells. Two days later, wells containing GFP positive cells were identified using an Evos M7000 automated microscope (4x objective). Supernatant from wells with GFP+ cells was harvested and a sequential series of limiting dilutions was performed to isolate GFP+ viruses. RNA was isolated from the supernatant of cells containing GFP+ viruses (Macherey-Nagel, #740956), reverse transcription reactions were done using SuperScript VILO Master Mix (Invitrogen, #11755050), and the GFP region containing the frameshift was amplified with Primestar GXL using gene specific primers (**Supplementary table 5**). The amplicon was gel purified (Macherey-Nagel, #740609) and its sequence determined using Sanger (Genewiz) or Oxford nanopore (Plasmidsaurus) sequencing.

### Generation of purified virions for deep RNA sequencing

For HCoV-OC43/fsGFP and HCoV-OC43/fsGFP(δNSP15) populations, cells (25 million 293T cells in 15cm dishes) were infected at an MOI of 2 PFU/cell with the P1 virus product of a CPER reaction transfection, and supernatant was harvested at 24h post-infection. Cell free supernatant (20mL) containing 8.2 x10^8^ PFU (HCoV-OC43/fsGFP(WT)) or 2.4 x10^8^ PFU (HCoV-OC43/fsGFP(δNSP15)) was clarified by centrifugation (1500 g for 10 minutes), and filtration (0.22µm). Virions were purified by centrifugation through a 30% sucrose cushion (in PBS) in a Beckman Optima XE-90 ultracentrifuge (Beckman SW32Ti rotor, 25,000 rpm, 4°C, 90 minutes). The virion pellet was resuspended in TRIzol reagent (Invitrogen, #15596026) and RNA was isolated according to the manufacturer’s protocol. The quality, integrity, and quantity of the resulting viral RNA was determined by Nanodrop measurement, agarose gel electrophoresis, and Qubit analysis, respectively. For SARS-CoV-2, 293T-ACE2 cells were transfected with WT SARS-CoV-2 CPER reaction products, and co-cultured with VeroE6 cells until cytopathic effect was observed. Supernatent (P1) from that coculture was applied to VeroE6 cells (5 million in a 10cm dish) that were cultured until cytopathic effect was observed (P2). This virus stock was expanded once more (P3; 50-75 million Vero cells in each of two 825cm 5 layer flasks). Virions were purified from a 20ml sample of supernatant which contained 3.6 x10^7^ PFU and virion RNA was isolated as described above for HCoV-OC43/fsGFP.

### Sequencing of virion RNA

Total RNA (300 ng) extracted from purified HCoV-OC43 and SARS-CoV-2 virions was used to generate RNA-Seq libraries using an Illumina Stranded Total RNA Prep kit (#20040529) and IDT xGen Stubby Adaptor _UDI Primers for Element (#10017037) following manufacture’s protocols. Libraries prepared with unique barcodes were pooled at equal molar ratios. The pool was denatured and sequenced on Element AVITI sequencer using AVITIOS 3.3.2 control software and Cloudbreak FS reagents to generate 2 x 150 nucleotide paired end reads, following manufacturers protocol (Document #MA-00008 Rev. K). Bases2fastq version: 2.4.0.2499042781 was used for adaptor removal and demultiplexing. The quality of the resulting reads was assessed using FastQC (v0.12.1). Reads were processed with Trimmomatic (v0.40) with the following settings: HEADCROP:10 SLIDINGWINDOW:4:25, paired-end mode, phred33, and genomes were assembled using Spades (v4.2.0) for HCoV-OC43/fsGFP(WT) and HCoV-OC43/fsGFP(δNSP15). Reads were aligned to the assembled reference genomes using the mem function of BWA-mem2 (v2.3) with default parameters. SARS-CoV-2 reads were processed as for HCoV-OC43 reads except de novo genome assembly was not performed and alignment was done against the Wuhan-1 reference genome (NCBI Reference Sequence: NC_045512.2). For Tiled-ClickSeq data, primer sequences were first removed as previously described^28^. Coverage maps were created with the genomecov function of bedtools (2.31.1) and plotted using Graphpad Prism.

### Infection of mice with SARS-CoV-2 and generation of infected mouse lung RNA

Female K18-hACE2 transgenic mice (B6.Cg-Tg(K18-ACE2)2Prlmn/J, strain #034860), were obtained from The Jackson Laboratory. Mice were 10-12 weeks old at the start of experiments and were housed at 22 °C with 30–70% humidity on a 12 h light/12 h dark cycle, with *ad libitum* access to food and water; they were acclimatized for at least 2 weeks before experimental procedures. All animal procedures were conducted in accordance with protocols approved by the Rockefeller University Institutional Animal Care and Use Committee (Protocol #24016-H).

Mice were anesthetized with inhalant Isoflurane administered at 3-4% in oxygen, and intranasally challenged with 2x10⁴ PFU of SARS-CoV-2/δORF7/GFP in a total volume of 30 µL. At 72 hours post-infection, mice were euthanized by CO2 inhalation in an isolated chamber with compressed carbon dioxide gas, and confirmation of respiratory arrest was followed by cervical dislocation to confirm death. Lungs were harvested immediately and stored at –80C until processing for total RNA extraction. Left and right lung lobes were processed separately using a Trizol based RNA extraction. SARS-CoV-2 genomic RNA was quantified by RT-qPCR targeting the N gene using the Luna® Universal qPCR Master Mix kit obtained from New England Biolabs. The primers used were Integrated DNA Technologies nCOV_N1 Forward Primer Aliquot, 50nmol, catalog #10006821, and Integrated DNA Technologies nCOV_N1 Reverse Primer Aliquot, 50nmol, catalog #10006822. Viral RNA copies determined with the aid of Integrated DNA Technologies 2019-nCOV_N_Positive Control standard, catalog #10006625. An RT-qPCR reaction for a housekeeping gene (GAPDH) was run in parallel to ensure comparable RNA concentrations among reactions.

### Generation of amplicons for sequencing

For purified SARS-CoV-2 virions, cDNA was synthesized with Superscript VILO Mastermix. For infected mouse lung tissue RNA cDNA was synthesized using Superscript III reverse transcriptase (Thermo-Fisher, #18080044) and a gene specific primer (**Supplementary table 5**) due to the high complexity of RNA in the sample. Amplicons (162 base pairs, including PCR primer sequences) were generated using the above cDNA templates and primers flanking the SARS-CoV-2 Spike furin cleavage site (**Supplementary table 5**) using Primestar GXL DNA Polymerase and purified by gel extraction. Prior to sequencing, DNA amplicons originating from two different infected mouse lung RNA samples were pooled at equimolar ratios.

### Sequencing of DNA amplicons

PCR amplicons (10 ng) were used to generate libraries using Illumina TruSeq Nano DNA Low Throughput Library Prep kit (#20015964), following manufacturers protocol. Libraries prepared with unique dual indexes were pooled at equal molar ratios. The pool was sequenced on Illumina NextSeq 2000 P4 flowcell using NextSeq Control Software v1.7.1.46395 to generate 150 nucleotide paired reads, following manufactures protocol (Document #200027171 v04). Bcl2fastq v2.20.0.422 was used for the adaptor removal and demultiplexing. PCR primers were removed from the resulting sequencing reads using Cutadapt (v5.2) with the following settings: -j 4 -g “GACATACCCATTGGTGCAGG…ctaataactctattgccatacccaca;optional” -g “tgtgggtatggcaatagagttattag…CCTGCACCAATGGGTATGTC;optional” -G “tgtgggtatggcaatagagttattag…CCTGCACCAATGGGTATGTC;optional” -G “GACATACCCATTGGTGCAGG…ctaataactctattgccatacccaca;optional” --discard-untrimmed. Reads were further quality filtered with Trimmomatic (MINLEN: 100, SLIDINGWINDOW4:25). BWA-mem2 mem was used to align reads to the SARS-CoV-2 (Wu-1, NCBI Reference Sequence: NC_045512.2) genome. A python program (tandem_dup_check.py) was used prior to analysis to identify and remove all tandem-duplicate insertions.

### Processing of RNA-sequencing data from clinical datasets

RNA-sequencing files were downloaded from the NCBI Sequence Read Archive (PRJNA805055, PRJNA613958). Quality was assayed with FastQC and bases were trimmed using the following Trimmomatic settings: phred33, LEADING:20 TRAILING:20 SLIDINGWINDOW:4:20. Reads were then aligned to SARS-CoV-2 Wuhan-1 (PRJNA613958 - NCBI Reference Sequence: NC_045512.2) or SARS-CoV-2 BA.1 (PRJNA805055 - NCBI Reference Sequence: OL672836.1) using BWA-mem2. Aligned reads were sorted, indexed (Samtools) and primer masking (PRJNA613958: ARTIC v3; PRJNA805055: ARTIC v4.1) was performed with iVar (v1.4.4) trim using the following settings: -m 30 -q 20 -s 4 -e.

### Insertion mutation detection and mapping

Inserted sequences of 2 nuclotides or greater were detected using a custom Python program using the Pysam (v0.23.3) software package (getins.py). CIGAR strings were used to detect the genome coordinates (+0) and sequence of inserts. Each sequence insertion was mapped back to the cognate reference genome, and the genomic coordinates of the insert origins were recorded. (+1, ins_origins.py) The distance between insertion origin and insertion destination was calculated as the absolute value of the difference between their genetic coordinates. To detect chimeric inserts, the 5’ end of inserts was mapped to the viral genome until a mismatch was detected. Next, the remaining sequence was mapped from that position. Only sequences which mapped completely to the viral genome in two fragments (without any mismatches) were designated as chimeras (find_chimeras.py).

### Designation of inserts of probable viral origin

To calculate the probability of an insert matching the viral genome by chance, every possible k-mer of a given length was extracted from both the + and – sense viral genome RNA using a custom python program (kmer.py). To allow for mismatches, for each k-mer extracted from the genome, sequences with every possible mismatch (up to the number specified) were calculated. The probability of a k-mer of a given length K occurring in the viral genome was calculated as the total number of unique k-mers generated from the genome divided by the total number of possible k-mers of that length (4^k^). Inserts with a <5% chance of matching a viral k-mer were designated of probable viral origin: 11+ nucleotides with 0 mismatches, 13+ nucleotides with 1 mismatch, 16+ nucleotides with 2 mismatches, and 18+ nucleotides with 3 mismatches.

### Dinucleotide junction analysis

For this analysis, inserts with ambiguous destination sites were excluded (ambiguous_sites.py) because their sequence identity could not be known with certainty (see **Extended Data Fig. 3**). For each insertion, the junction dinucleotides were defined as the [last nucleotide of the upstream segment][first nucleotide of the downstream segment] at both the 5′ and 3′ boundaries of the insert origin site. Monte Carlo permutation testing was used to produce null values for each of the 16 dinucleotide frequencies: For each insertion sequence, the dinucleotide at a random coordinate in the genome was recorded. The average dinucleotide frequencies from 10,000 permutations of this analysis were defined as the null values. Fold enrichment (Obs/Exp) was calculated as the experimentally calculated dinucleotide frequencies divided by the null value for each dinucleotide. Bootstrap analysis (500 replicates) was used to calculate a standard error of the mean (SEM) (dinuc_origins.py) The same process was repeated for the insert destination site, where junction dinucleotides were defined as last nucleotide of the genome sequence immediately 5’ to the insert followed by the first nucleotide immediately 3’ to the insert in the positive sense viral genome. (dinuc_destination.py)

### Genomic distribution histograms and Circos plots

Plots were generated with custom Python programs (sliding_window_his_batch.py, circo_chim.py) using NumPy (v2.4.3) and matplotlib (3.10.9).

### Analysis of modifications of viral coding sequences by insertions

A custom Python program (rxxr_finder.py) and a virus specific BED file of genomic coordinates were used to map insertion sites to CDS (coding sequence) features. For analysis of potential furin cleavage site acquisition, insertions occurring outside of any CDS or causing a frameshift were excluded. For the remaining insertions, the modified CDS sequence was translated and inserts creating a stop codon were excluded. Finally, insertion events that created new RXXR motifs were tabulated.

## Data Availability statement

The sequencing datasets generated during the current study have been deposited in the NCBI Sequence Read Archive under BioProject accession number **PRJNA1509607.** The accession numbers for the publicly available data are listed in **Supplementary Table 4**.

## Code Availability

The custom Python scripts used for data analysis are publicly available on Github at https://github.com/cbianco1/coronavirus_insert_analysis.

## Acknowledgments

We thank the Rockefeller University Genome Resource center for technical advice and perfoming all sequencing in this study. PDB is a HHMI investigator. This article is subject to HHMI’s Open Access to Publications policy. HHMI lab heads have previously granted a non-exclusive CC BY 4.0 license to the public and a sublicensable license to HHMI in their research articles. Pursuant to those licenses, the author-accepted manuscript of this article can be made freely available under a CC BY 4.0 license immediately upon publication.

## Author contributions

CB and PDB conceived the work. CB, ML, MA, and MT performed the experiments or constructed reagents. CB and AS analysed the data, PDB and TH supervised the work CB, TH and PDB wrote the paper

## Supplementary Information

**Supplementary Table 1:** Quantification and analysis of GFP+ virions isolated from HCoV-OC43/fsGFP(WT) and HCoV-OC43/fsGFP(δNSP15) passage

**Supplementary Table 2:** Analysis of HCoV-OC43 derived kmers for designation of inserst as probable viral origin

**Supplementary Table 3:** Overall quantification of insert detection and origin

**Supplementary Table 4** Accession numbers for publicly available datasets used herein

**Supplementary Table 5** Oligonucleotides used for cloning and amplicon generation

**Supplementary Dataset 1:** Catalogs of inserts from HCoV-OC43/fsGFP(WT) and HCoV-OC43/fsGFP(δNSP15) found in purified virion sequencing libraries

**Supplementary Dataset 2:** Catalogs of inserts from SARS-CoV-2(WT) and SARS-CoV-2(δNSP15) found in published sequencing datasets

**Supplementary Dataset 3:** Catalogs of inserts from SARS-CoV-2 found in purified virion sequencing libraries

**Supplementary Dataset 4:** Catalogs of inserts from SARS-CoV-2 found in amplicon sequencing libraries

**Supplementary dataset 5:** Catalogs of inserts from SARS-CoV-2 found clinical datasets

For supplementary datasets 1-5, the genomic coordinates of insert origins and insert destinations are denoted by ‘position’ and ‘ref_coord_before_insertion’, respectively. In the position column, ‘duplicate’ and ‘NA’ indicate that the insertion sequence occurs more than once or not at all in the viral genome, respectively. The ‘mismatches’ column states the number of mismatches between the insert and its closest match in the viral genome (with more than 4 mismatches reported as no_match). The ‘strand_mapping’ column in the sheets with the suffix ‘_inserts’ indicates the viral genome strand that the insert originates from. The full sequencing read, the insert sequence, and the length of the insertion (nt) are denoted by ‘full_sequence’, ‘insert_sequence’, and ‘insert_length’, respectively. The tables containing the suffix ‘_coding’ contain information on the coding potential of inserts in the context of their destination site. In the column ‘creates_rxxr’, insertions occurring outside of a CDS (coding region) are labeled ‘N/A’ while inserts that are in a coding region are labeled ‘stop_codon_introduced’, ‘not_in_frame’, no, and yes to indicate if they introduce a stop codon, create a frameshift, are in frame but do not create a new RXXR motif, or are in frame and do create a new RXXR motif, respectively. ‘Annotation’ states which CDS the insertion occurs in, if any. ‘rxxr_motif’ lists the amino acid sequence of new RXXR motif(s) created by the insertion. To remove redundant insertion events, insertions were deduplicated by identical (‘insert_sequence’, ‘ref_coord_before_insertion’) pairs, followed by deduplication by ‘read_id’.

---

**Extended Data Fig. 1.**
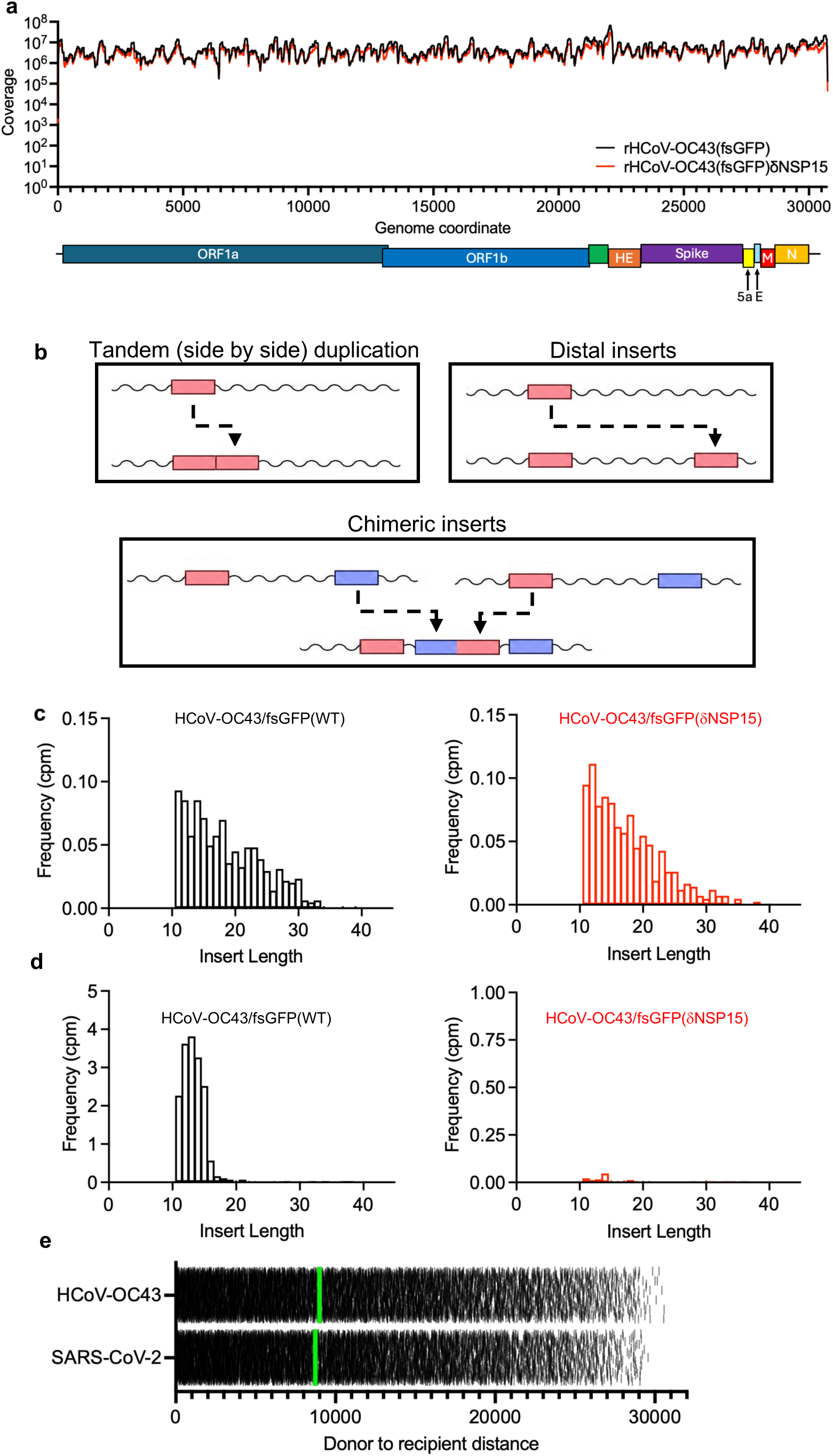
Analysis of HCoV-OC43 virion RNA sequencing data. (a) Overall coverage in the rHCoV-OC43/fsGFP RNA seq experiment. Coverage (Y-axis) indicates the number of sequence reads for each nucleotide position in the rHCoV-OC43/fsGFP(WT) (black) and rHCoV-OC43/fsGFP(δNSP15) (red) genomes. (b) Schematic representation of the different classes of insert mutations described herein. (c) Frequency distribution of insert lengths (in nucleotides) for tandem duplications (side by side inserts) of 11 nucleotides or longer in sequencing reads, given as counts per million sequencing reads (cpm), from RNAseq analysis of purified HCoV-OC43/fsGFP virions WT (left, black) and δNSP15 (right, red). (d) Frequency distribution of insert lengths (in nucleotides) for distal inserts of probable viral origin in sequencing reads, given as counts per million sequencing reads (cpm), from RNAseq analysis of purified HCoV-OC43/fsGFP virions WT (left, black) and δNSP15 (right, red). (e) Distances (in nucleotides) between randomly generated sites in the HCoV-OC43 and SARS-CoV-2 genomes. Each tick represents a different pair of random sites, green lines = median distance.

**Extended Data Fig. 2.**
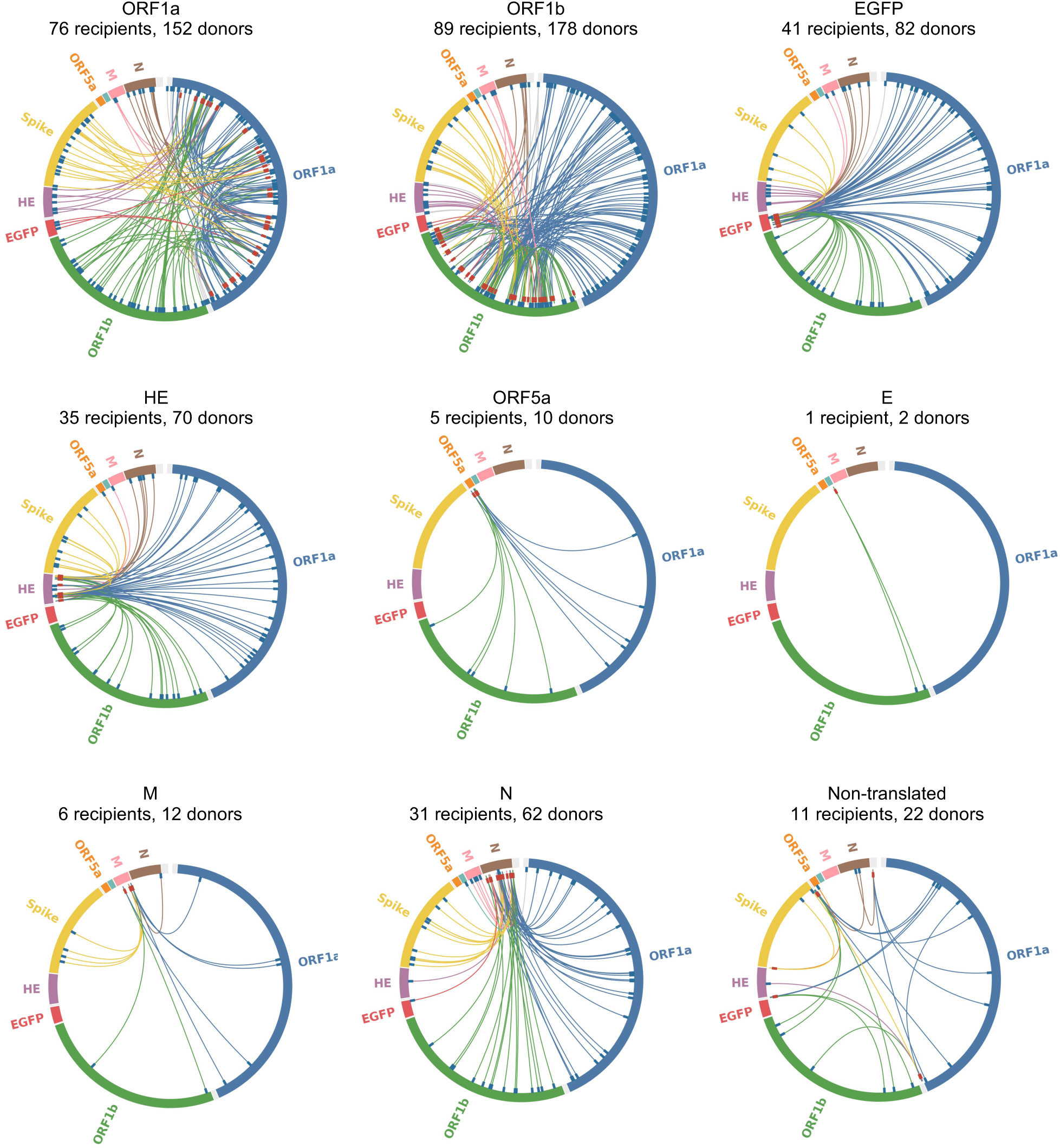
Chimeric inserts in HCoV-OC43 RNA genomes. Circos plots, indicating locations of donor sequences (blue ticks, outer circle) and recipient sites (red ticks, inner circle) for chimeric inserts with two viral insert donors into each recipient site in each of the open reading frames and non-translated sequences of rHCoV-OC43/fsGFP(WT).

**Extended Data Fig. 3.**
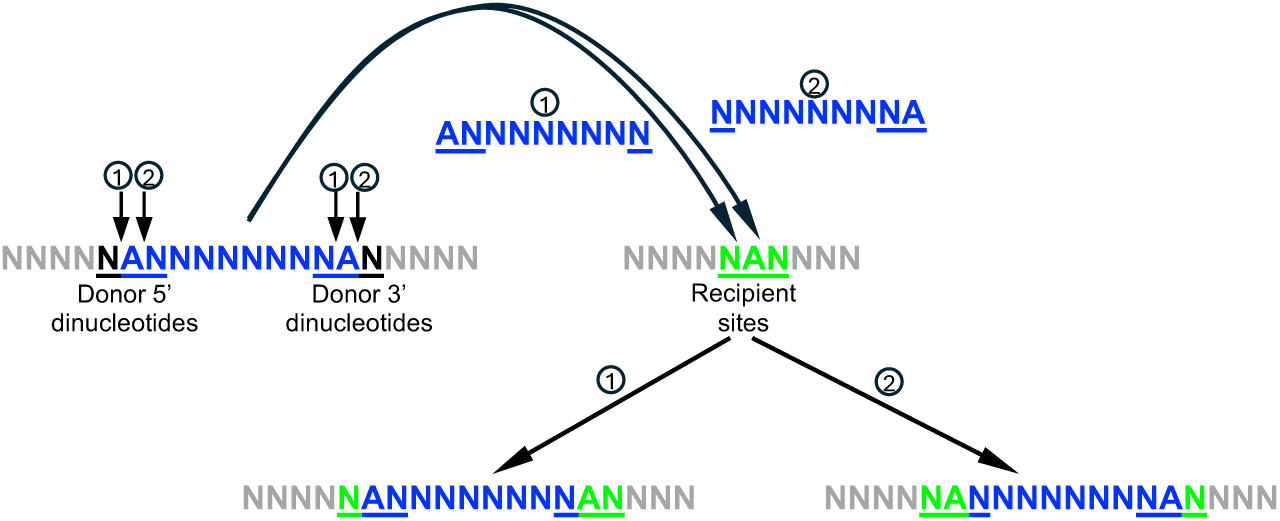
Ambiguity in some insert junctions. (d) Schematic illustrating the source of ambiguity in 5’ and 3’ donor dinucleotides (blue/black) and recipient dinucleotides (green), when the same base (in this case A) is present at the ends of the insert and in the recipient site. Inserts that exhibited this ambiguity were excluded from the sets that were analyzed for donor and recipient dinucleotide compositional bias.

**Extended Data Fig. 4.**
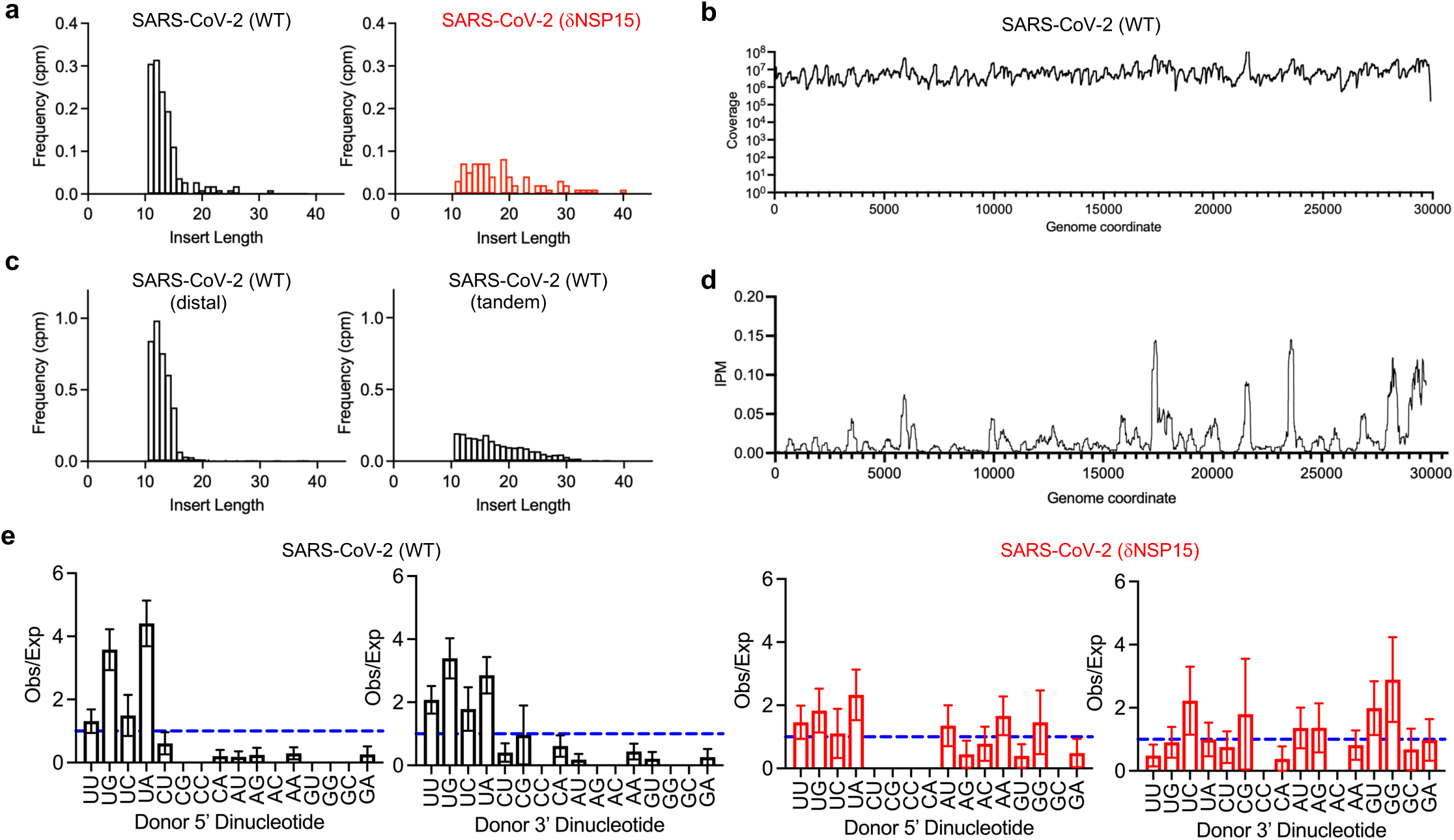
Analysis of SARS-CoV-2 virion RNA sequencing data. (a) Frequency distribution of insert lengths (in nucleotides) for distal inserts of probable viral origin in sequencing reads, given as counts per million sequencing reads (cpm), from published sequencing datasets (Zhou et al) for SARS-CoV-2(WT) (left, black) and SARS-CoV-2(δNSP15) (right, red). (b) Overall coverage in the SARS-CoV-2 RNA seq experiment done herein. Coverage (Y-axis) indicates the number of sequence reads for each nucleotide position in the SARS-CoV-2 genome. (c) Frequency distribution of insert lengths (in nucleotides) for distal inserts of probable viral origin (left) and tandem (side-by side) duplications (right), given as counts per million sequencing reads (cpm), in the SARS-CoV-2 virion RNAseq dataset. (d) Frequency of tandem (side-by-side) duplications in a sliding window of 250 nucleotides across the SARS-CoV-2 genome, given as inserts per 250 nucleotides per million reads (IPM) in the sequenced virion population. (e) Observed/expected (Obs/Exp) frequency of the occurrence of dinucleotides at the 5’ and 3’ ends of inserts in the context of unambiguous insert donor sites for distal inserts of probable viral origin in published (Zhou at al) SARS-CoV-2(WT) (left, black) and SARS-CoV-2(δNSP15) (right, red) datasets.

**Extended Data Fig 5.**
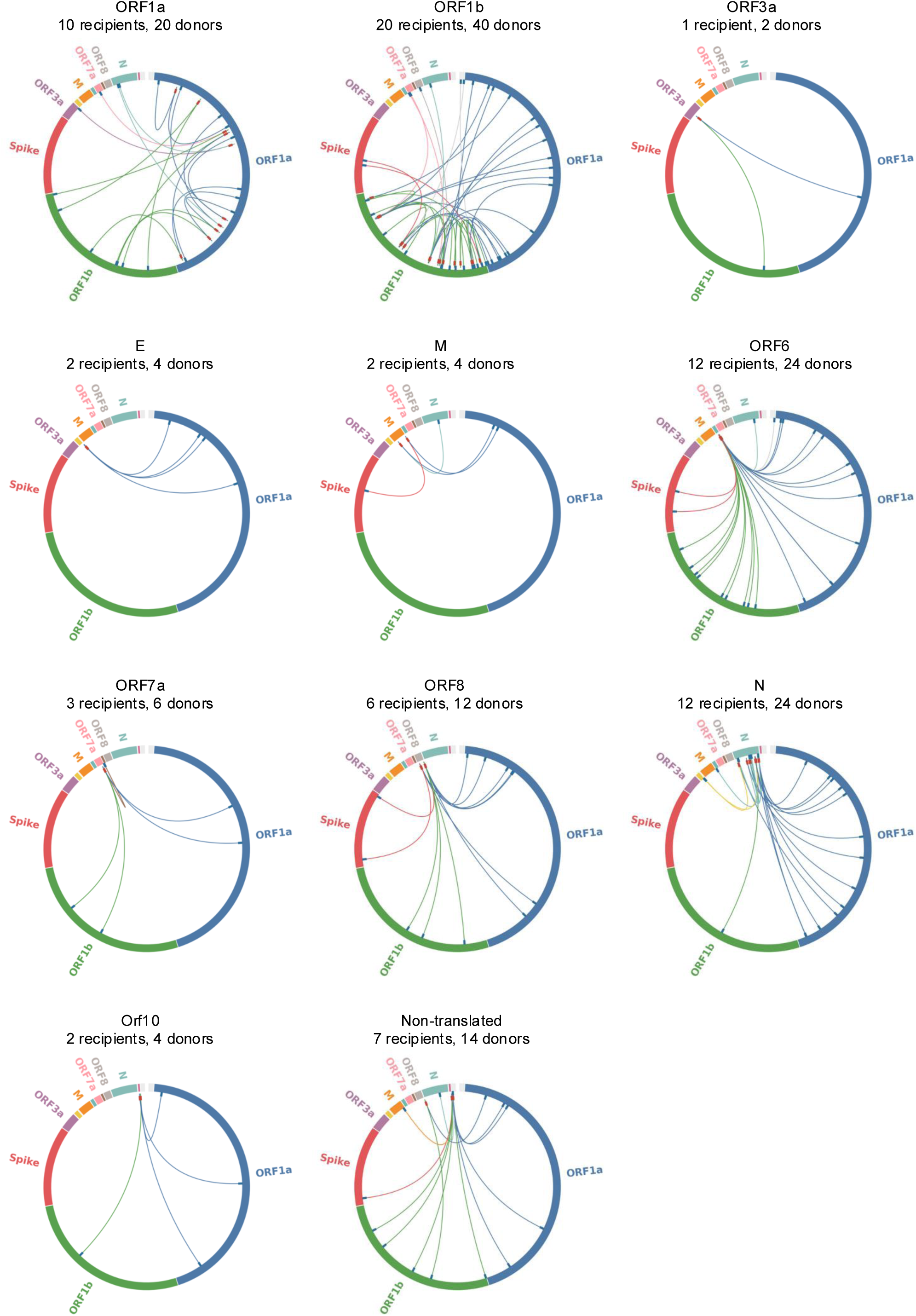
Chimeric inserts in SARS-CoV-2 RNA genomes. Circos plots, indicating locations of donor sequences (blue ticks, outer circle) and recipient sites (red ticks, inner circle) for chimeric inserts with two viral insert donors into each recipient site, in each of the open reading frames and non-translated sequences of SARS-CoV-2.

**Extended Data Fig 6.**
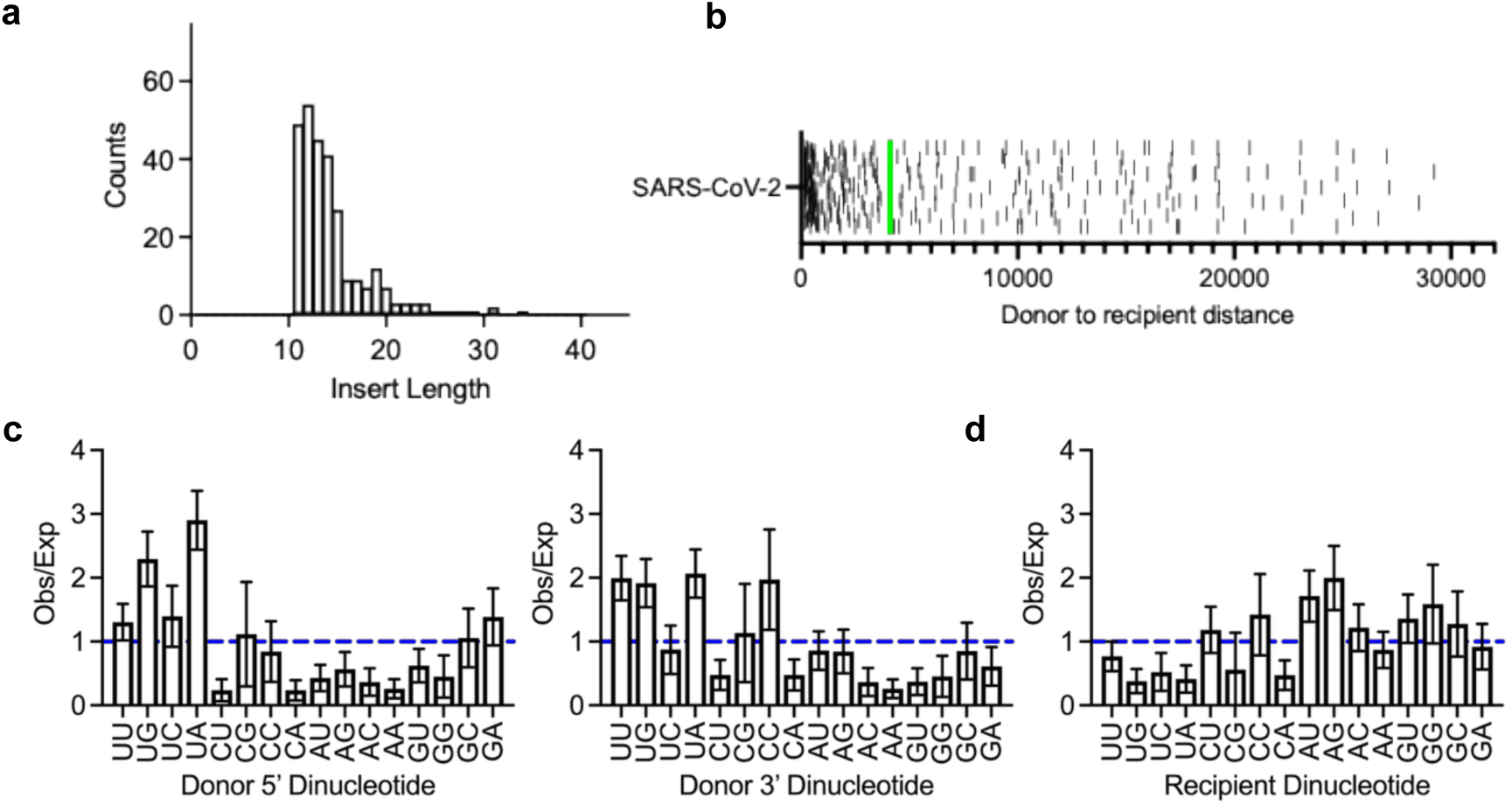
Analysis of SARS-CoV-2 insertional mutants in a clinical datset. (a) Frequency distribution of insert lengths (in nucleotides) for distal inserts of probable viral origin of 11+ nucleotides in sequencing reads, from a published sequencing dataset (PRJNA805055) from SARS-CoV-2 infected humans, given as raw counts. (b) Distances (in nucleotides) between donor and recipient sites for distal inserts of probable viral origin for in a published sequencing dataset (PRJNA805055) from SARS-CoV-2 infected humans. Each tick represents a different insertion event, green lines = median distance. (c, d) Observed/expected (Obs/Exp) frequency of the occurrence of dinucleotides at the 5’ and 3’ ends of inserts in the context of unambiguous insert donor sites (c) and recipient sites (d) for distal inserts of probable viral origin from a published sequencing dataset (PRJNA805055) from SARS-CoV-2 infected humans.

**Extended Data Fig 7.**
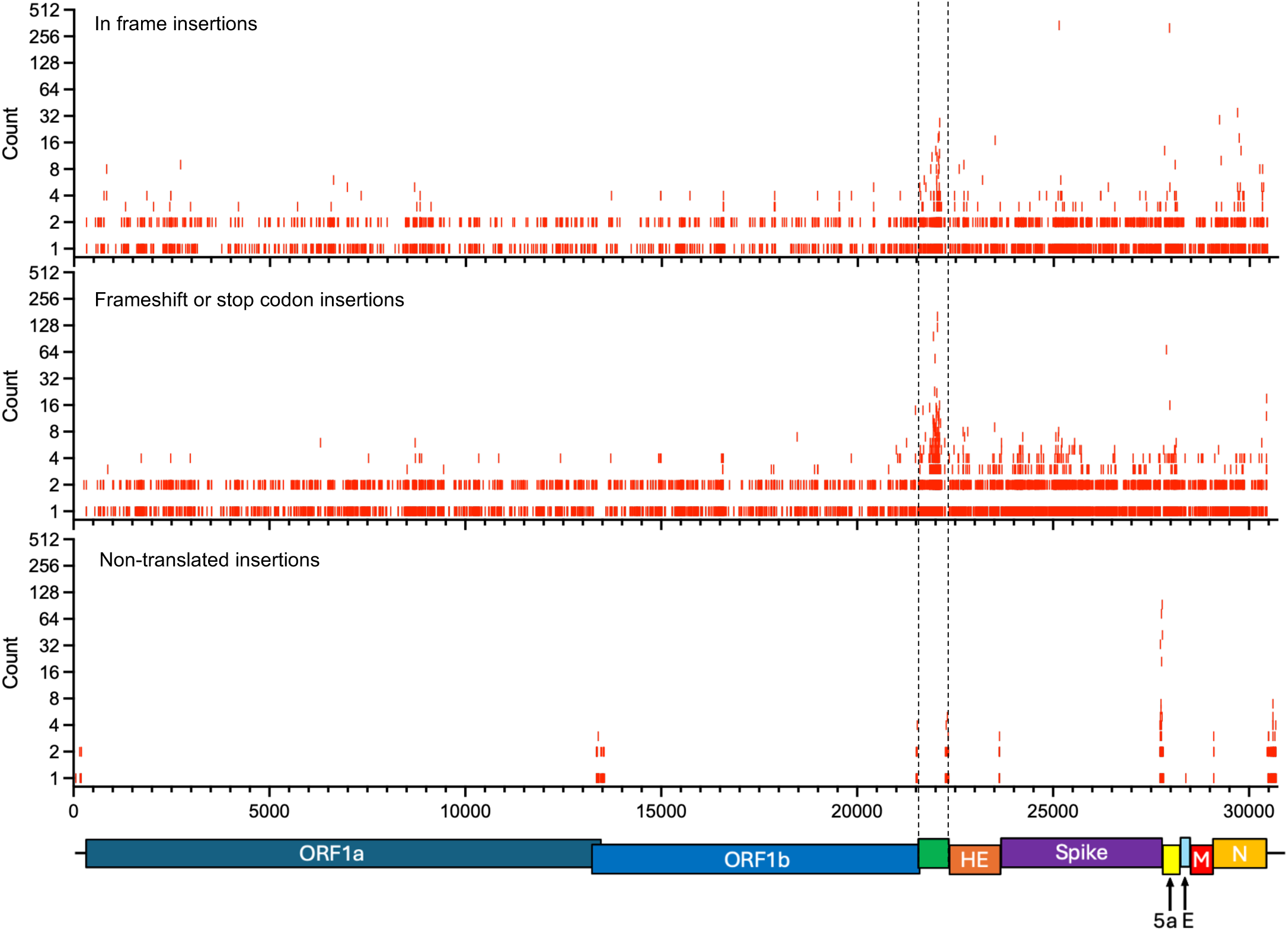
Distribution, frequency and coding potential of insertion mutations in HCoV-OC43/fsGFP virion RNA. Charts, with the HCoV-OC43/fsGFP genome represented on the X-axis showing the number of reads (out of 644,501,025 total mapped reads) in which individual insertional mutants in open reading frames were present in the sequenced virion population that generate in frame insertions (upper), protein truncations (middle), or occurred in untranslated regions (lower). Each individual red tick represents an insertion event. Dashed vertical lines indicate the boundaries of the inserted eGFP reporter gene.

**Extended Data Fig 8.**
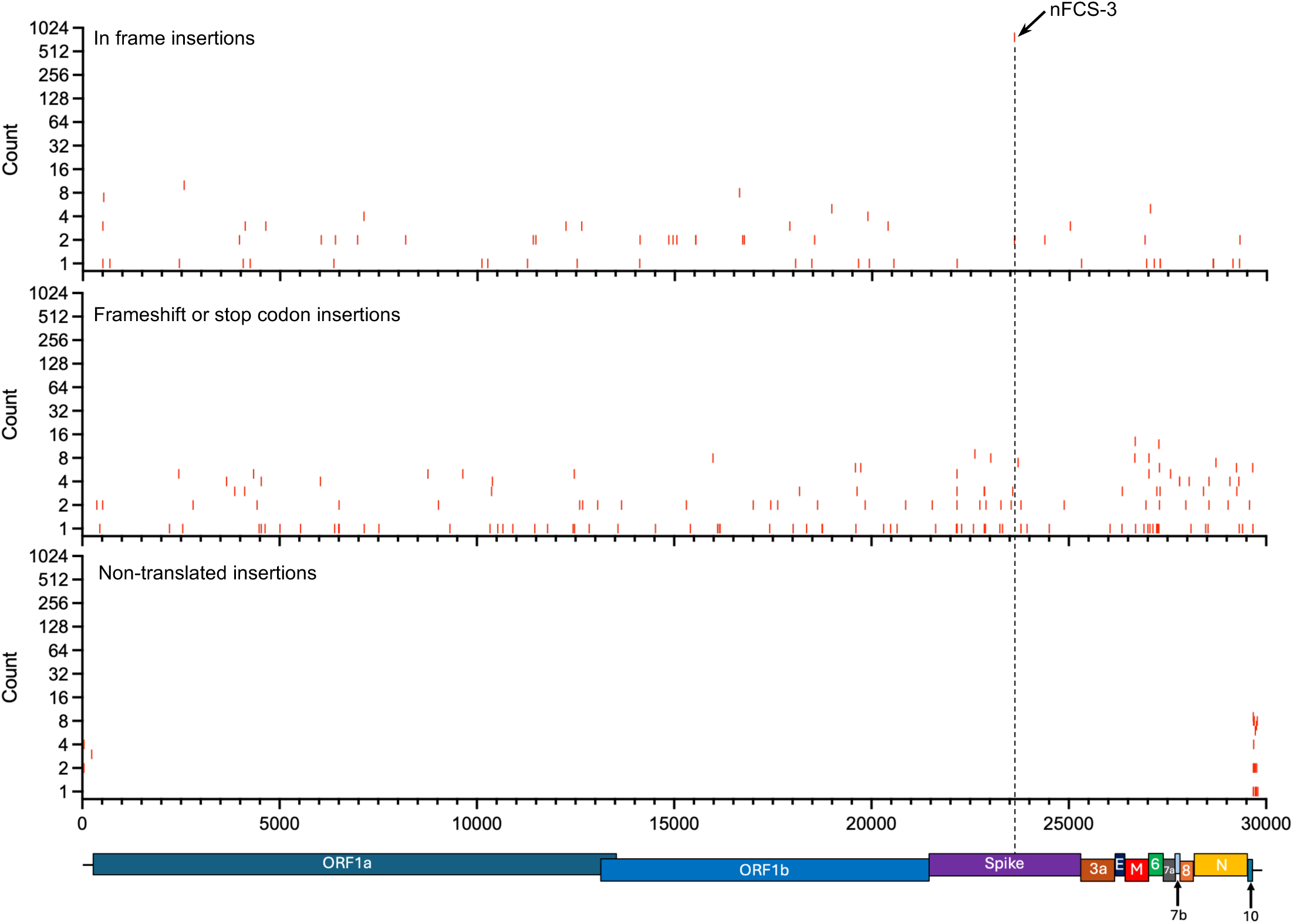
Distribution, frequency and coding potential of insertion mutations in SARS-CoV-2 virion RNA. Charts with the SARS-CoV-2 genome position represented on the X-axis showing the number of reads (out of 108,408,396 total mapped reads) in which individual insertional mutants in open reading frames were present in a published sequencing dataset (Zhou et al) that generate in frame insertions (upper), protein truncations (middle), or occurred in untranslated regions (lower). The datapoint corresponded to the appearance of a new furin cleavage site (nFCS-3), at the site of the existing SARS-CoV-2 spike furin cleavage site (vertical dashed line) is highlighted.

**Extended Data Fig 9.**
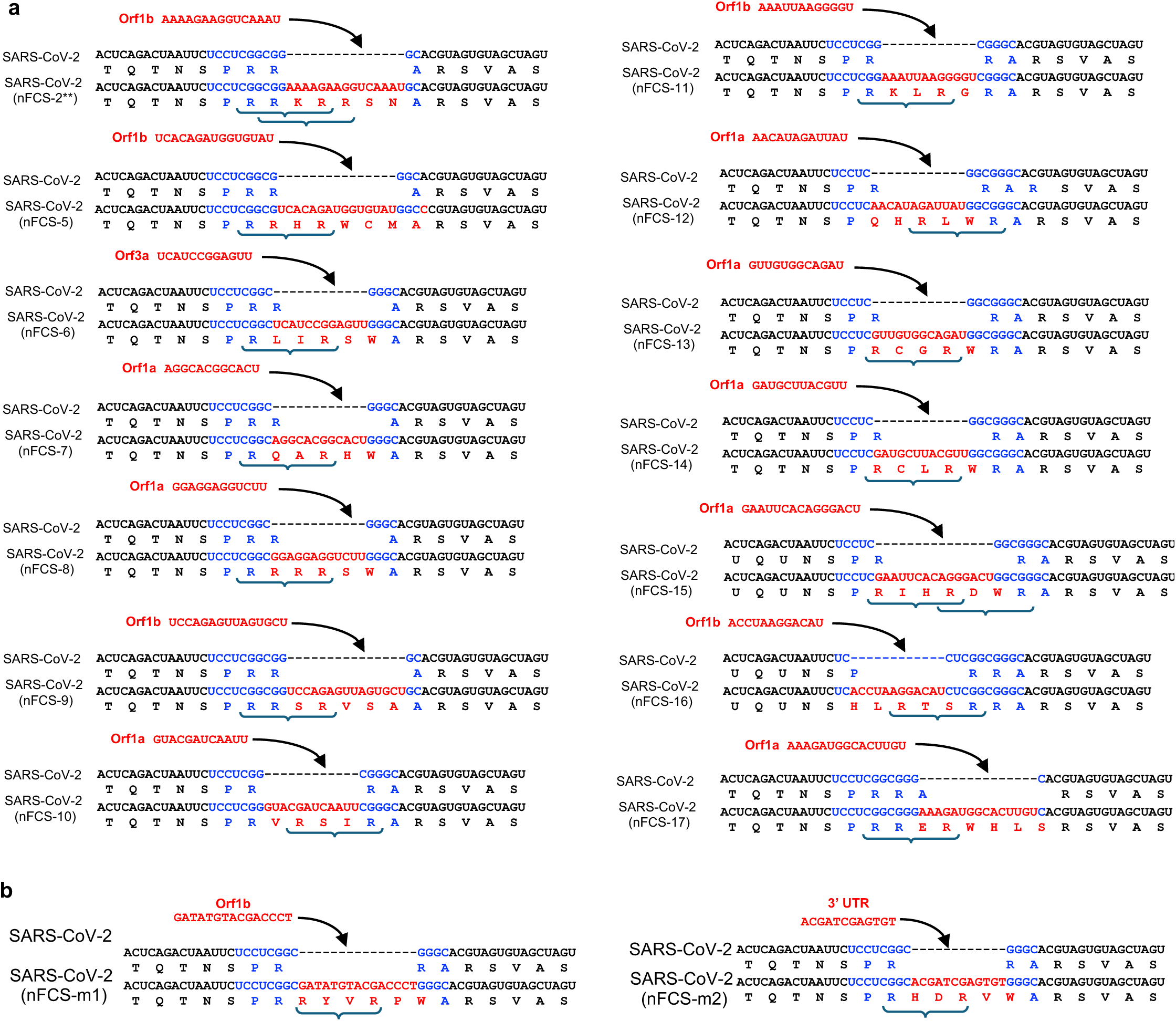
Genesis of new or replacement furin cleavage sites during SARS-CoV-2 replication. (a, b) Additional examples of acquisition of novel furin cleavage sites at the site of the existing SARS-CoV-2 furin cleavage site during the course of SARS-CoV-2 replication. The pre-exisiting SARS-CoV-2 FCS is shown in blue, sites acquired during replication of molecularly cloned virus are shown in red. Potential furin cleavage sites (RXXR motifs) are bracketed. The nFCS-2 was found in both the purified virion RNA sequencing dataset, and the virion-derived PCR amplicon dataset, while nFCS-5 through 17 (a) were found only in the virion-derived PCR amplicon dataset. The nFCS-m1 and nFCS-m2 (b) were found only in the infected mouse lung amplicon dataset.

**Extended Data Fig 10.**
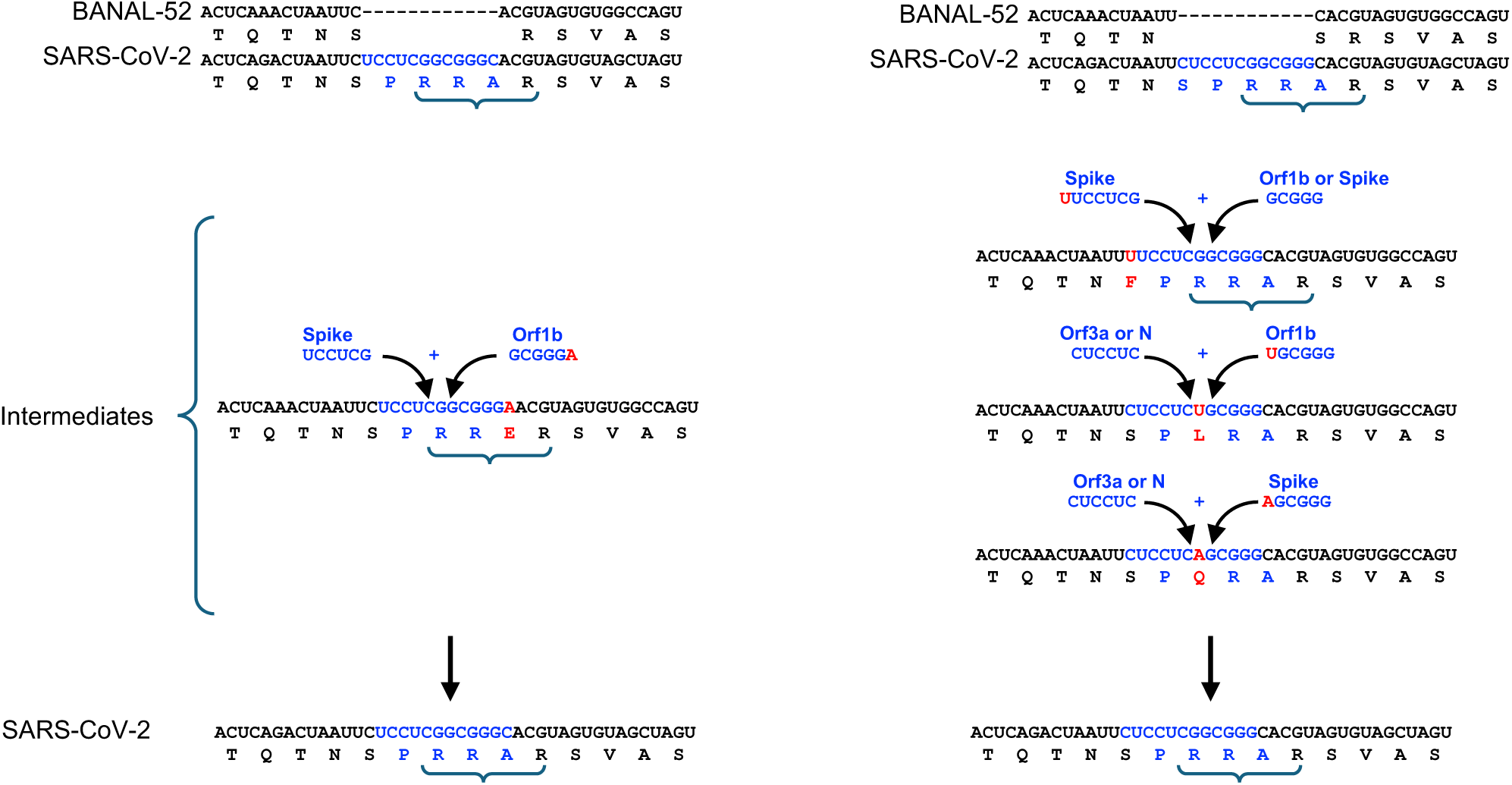
Potential models for the genesis of the SARS-CoV-2 furin cleavage site. Alternative models for the origin of SARS-CoV-2 FCS. Black text represents the sequence of BANAL20-52, a close relative of SARS-CoV-2 found in *Rhinolophus* bats. Blue text represents two possible insert configurations (which is ambiguous based on alignment of the BANAL20-52, and SARS-CoV-2 genomes). Several potential alternative insertion events that could generate the SARS-CoV-2 prototype sequence are depicted. Red letters indicate mismatches to the SARS-CoV-2 (WT) sequence. Potential furin cleavage sites (RXXR motifs) are bracketed.

